# A “non-muscle” α-actinin is an intrinsic component of the cardiac Z-disc and regulates sarcomere turnover, contractility, and heart remodeling

**DOI:** 10.1101/2024.11.26.625523

**Authors:** James B Hayes, Dylan Ritter, Abigail C Neininger-Castro, Alaina H Willet, Leah R Caplan, Yu Wang, Xiao Liu, Nilay Taneja, Zachary C Sanchez, Kyra Smart, Cynthia A Reinhart-King, Qi Liu, Matthew J Tyska, Erdem D Tabdanov, Quinn S Wells, Ela W Knapik, Dylan T Burnette

## Abstract

Cardiac sarcomeres generate the fundamental forces behind each heartbeat and are thought to contain only muscle-specific cytoskeletal proteins. We show that a widely expressed actin cross-linking protein, α-actinin 4 (ACTN4), is a sarcomere component of the human and zebrafish heart *in vivo* and in human iPSC-derived cardiac myocytes (CMs) *in vitro*. A confluence of biochemical experiments, immunofluorescence, and AI modeling suggest ACTN4 forms a heterodimeric complex with muscle-specific ACTN2 at the cardiac Z-disc, the cardiac sarcomere border. ACTN4 depletion from human iPSC-CMs stabilizes canonical sarcomere proteins and drives contractility-dependent cellular hypertrophy while ACTN4 overexpression destabilizes sarcomeres. ACTN4 depletion from zebrafish embryos specifically increases ventricular contractility which drives atrial enlargement, suggesting biomechanically driven atrial remodeling. ACTN4-associated phenotypes in both model systems lack hallmarks of cardiac disease models and an ACTN4 variant in humans is associated with reduced risk for disease. Our findings suggest a “non-muscle” actinin regulates heart contractility and influences clinical outcomes related to heart failure.

## INTRODUCTION

Cardiac muscle must produce sufficient force to propel blood through the circulation, which is its workload^1,2^. Changes in cardiac workload can stimulate structural remodeling of cardiac muscle^2–4^. Cardiac remodeling is directly coupled to workload by the cardiac sarcomere^2,5^, the fundamental unit of cardiac contractile force^6^. Cardiac sarcomeres generate contractile force primarily through cardiac muscle-specific proteins^7–9^. Mutations in cardiac sarcomere proteins that impact force generation can drive cardiac remodeling at the tissue-level^10–12^. In those mutations, whether contractile force is heightened or depressed determines the clinical course of remodeling and underlies conditions like heart failure with preserved ejection fraction (HFpEF)^13–15^. The association of remodeling with disease has centered most cardiac research around pathophysiological remodeling, even though remodeling is not inherently pathological.

Cardiac sarcomeres generate force as heart muscle-specific myosins bind and contract overlapping filaments of actin^6,16,17^. Force generation is counterbalanced at the borders of each sarcomere (i.e., Z-discs) where actin filaments of adjacent sarcomeres are anchored and mechanically linked by the muscle-specific actin cross-linker, α-actinin 2 (ACTN2)^8,18,19^. ACTN2, and other muscle-specific sarcomere proteins, represent specialized versions of otherwise common proteins that are expressed in a variety of cell types^20–22^. Those widely expressed proteins that have been studied in muscle have appeared only to function during sarcomere assembly^23,24^. This has led to a paradigm where widely expressed versions of muscle proteins, commonly called “non-muscle” proteins, serve only as temporary sarcomere building blocks^23–28^.

Here, we report that a widely-expressed paralog of ACTN2, ACTN4, is a component of the cardiac Z-disc in both the adult human and mouse heart and in developmental models of human and zebrafish. In developmental models, ACTN4 is non-essential for the basic assembly of the sarcomere lattice. Instead, it functions as a negative regulator of sarcomere stability, contractility, and cellular hypertrophy. Depletion of ACTN4 from cardiac myocytes reduces sarcomere protein turnover and leads to contractility-dependent hypertrophy. The increase in contractility appears to be phenotypically distinct from well-established models of cardiomyopathy. *In vivo,* ACTN4 depletion from zebrafish embryos increases ventricular ejection fraction which drives contractility-dependent atrial remodeling. Furthermore, computational genetics approaches through Vanderbilt’s biobank of de-identified patient records (BioVU) reveal certain single nucleotide polymorphisms associated with reduced expression of ACTN4 are associated with decreased risk of heart failure with preserved ejection fraction (HFpEF) at the population level, suggesting certain ACTN4 variants may be cardioprotective.

## RESULTS

### ACTN4 is a sarcomere protein in vivo and in iPSC derived cardiac myocytes

Our study began with data we independently curated from publicly available datasets of protein expression within the normal (non-diseased), adult human heart^29–31^. Within these datasets we focused on the beating muscle cells of the heart, the cardiac myocytes (CMs). We generated panels comparing the relative expression within CMs of muscle-specific proteins and their widely-expressed paralogs. While some of the interrogated widely-expressed proteins were not expressed by adult CMs, others were expressed at appreciable levels relative to their muscle-specific paralogs **(Fig. 1A)**. We were especially struck by the relative expression levels of the ACTN2 paralog, ACTN4, which were the highest among all of the widely-expressed paralogs of muscle proteins and were comparable even to expression of muscle-specific MYH6 (α-myosin II) **(Fig. 1A).** However, while MYH6 has been the subject of decades of study by the cardiovascular research community^32–34^, ACTN4 has gone completely unstudied and its impact, if any, on heart function is unknown.

**Figure 1:**
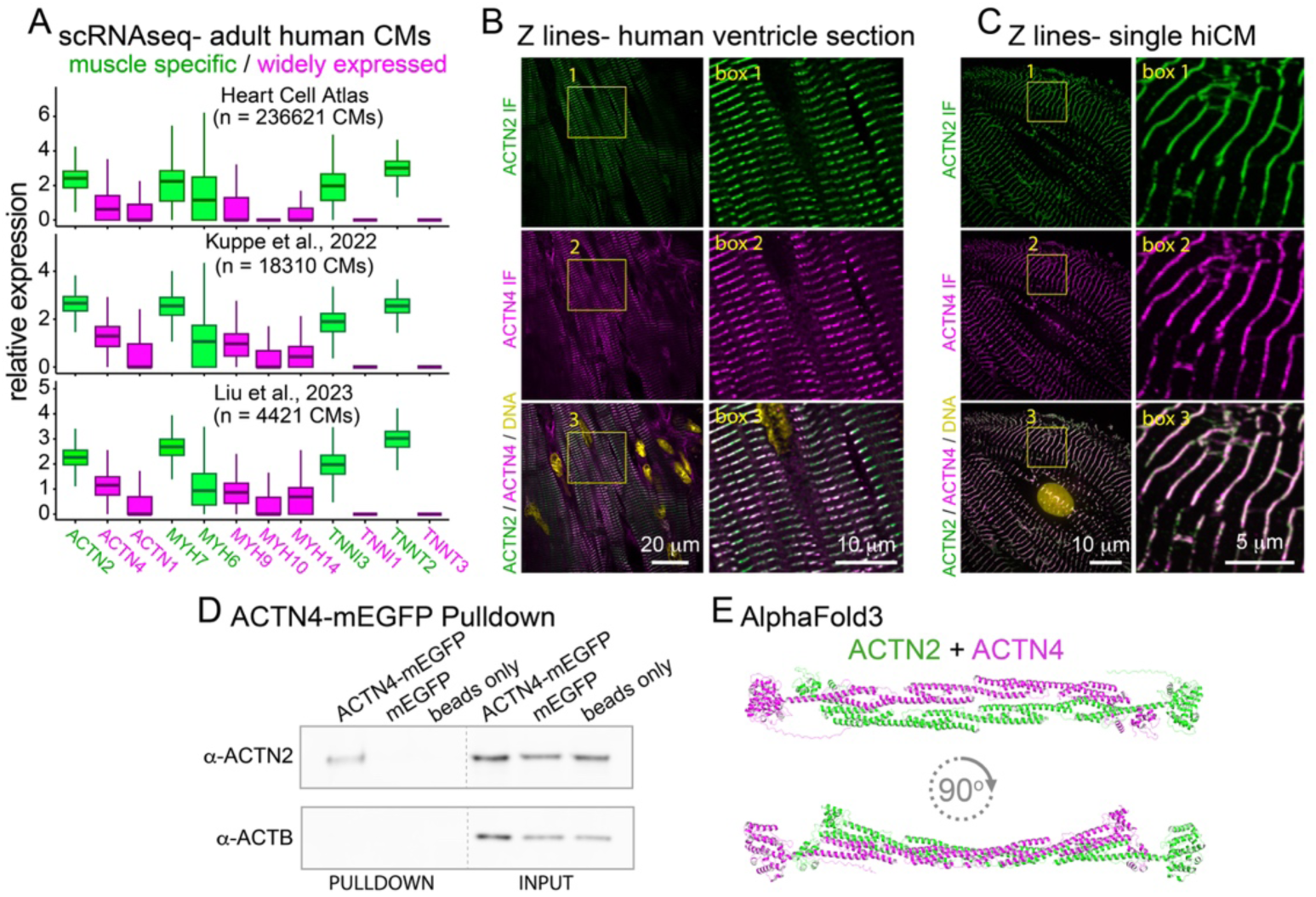
ACTN4 is a sarcomere protein in the adult heart and in iPSC-derived cardiac myocytes. **1A:** Relative expression levels of muscle-specific sarcomere proteins and their widely expressed paralogs in adult human cardiac myocytes. **1B:** Endogenous ACTN2 (green) and ACTN4 (magenta) in a section of non-diseased, adult human ventricle muscle. **1C:** Endogenous ACTN2 (green) and ACTN4 (magenta) in a human iPSC-derived cardiac myocyte (hiCM). **1D:** Co-immunoprecipitation of endogenous ACTN2 with ACTN4-mEGFP. **1E:** Predicted structure of an ACTN2:ACTN4 heterodimer by AlphaFold3.

To determine if we could localize ACTN4 within CMs, we identified a commercially available monoclonal antibody raised against the extreme ACTN4 N-terminus, which is unique among the actinin paralogs. Following validation using human iPSC-derived CMs (hiCMs) that the ACTN4 antibody did not bind non-specifically to ACTN2 or ACTN1 **(Fig. S1A-B),** we acquired formalin-fixed, paraffin-embedded samples of the adult, non-diseased human heart ventricle and mouse ventricle for immunofluorescence stains. ACTN4-specific antibodies localized within both tissues in a striated pattern **(Fig. S1C),** archetypical of muscle sarcomeres. Co-immunostains of ACTN4 and ACTN2 within the human ventricle revealed that ACTN4 striations were co-localized with ACTN2 striations **(Fig. 1B)**, indicating the sarcomere Z-disc. These data suggest ACTN4 is an evolutionarily-conserved sarcomere protein in the adult mammalian heart.

Because an unbiased, global proteomics study had identified ACTN2 and ACTN4 as proteins that could form a complex in HEK293 cells^35^, we hypothesized that ACTN4 and ACTN2 formed a complex at the CM Z-disc. To biochemically test our hypothesis, we transfected iPSC-derived human cardiac myocytes (hiCMs) with ACTN4-mEGFP, which localized to hiCM Z-lines (i.e., the two-dimensional projections of Z-discs) similar to immunofluorescence stains of endogenous ACTN2 and ACTN4 (**Fig. 1C, Fig. S1D**). Immunoprecipitation of ACTN4-mEGFP from lysed hiCMs using GFP-Trap beads revealed that endogenous ACTN2 co-immunoprecipitated with ACTN4-mEGFP **(Fig. 1D; Fig. S1E).** Actin did not precipitate **(Fig. 1D; Fig. S1E)**, suggesting we had pulled down ACTN4-mEGFP:ACTN2 complexes that were dissociated from actin filaments. To gain insight into the structure of a potential ACTN2:ACTN4 complex, we modeled an ACTN2:ACTN4 interaction *in silico* using the artificial intelligence tool AlphaFold3^36^. AlphaFold3 predicted ACTN2 and ACTN4 would form a heterodimer in a classical anti-parallel conformation, i.e., with actin-binding domains facing outward and dimerization facilitated by the two rod domains **(Fig. 1E)**, with similar confidence to the canonical ACTN2 homo-dimer **(Fig. S1F)**. Taken together, these data suggest a subset of actinin dimers at the cardiac myocyte Z-disc are anti-parallel ACTN4:ACTN2 heterodimers.

### ACTN4 is a negative regulator of sarcomere stability and contractility

Because ACTN4 and ACTN2 are in the same protein family and co-localized within CMs, we hypothesized they would have similar functions in CMs. ACTN2 is an essential sarcomere component^37,38^ and depletion of ACTN2 from hiCMs *in vitro* disrupts sarcomere assembly^25^. Therefore, we asked if ACTN4 depletion would similarly disrupt hiCM sarcomere assembly. Surprisingly, ACTN4-depleted hiCMs were enlarged and assembled more sarcomeres than either controls or ACTN4-depleted hiCMs ectopically expressing full-length ACTN4-mEGFP **(Fig. 2A-B; Fig. S2A-F)**. ACTN4 depletion did not impact ACTN2 levels **(Fig. S2C)**. Furthermore, ACTN4 overexpression in hiCMs caused a reduction in sarcomeres per cell that was not recapitulated by ACTN2 overexpression **(Fig. 2C),** suggesting ACTN4 and ACTN2 have distinct functions in CMs. Since α-actinin proteins function primarily as actin cross-linkers^39^, we next tested the hypothesis that the functions of ACTN2 and ACTN4 are distinguished by differences in actin cross-linking. Expression of the chimeric construct ACTN4ABD:ACTN2-mEGFP, which encodes the ACTN4 actin-binding domain (ABD) in frame with the remaining sequence of ACTN2, mimicked the dominant-negative effect of full-length ACTN4-mEGFP expression on the number of sarcomeres **(Fig. 2D)**. Conversely, expression of K255E-mEGFP, an ACTN4 point mutant with increased actin affinity^40^, increased the number of sarcomeres **(Fig. 2D)**. Taken together, these data suggest ACTN4 negatively regulates sarcomere number or stability through differences in actin cross-linking.

**Figure 2:**
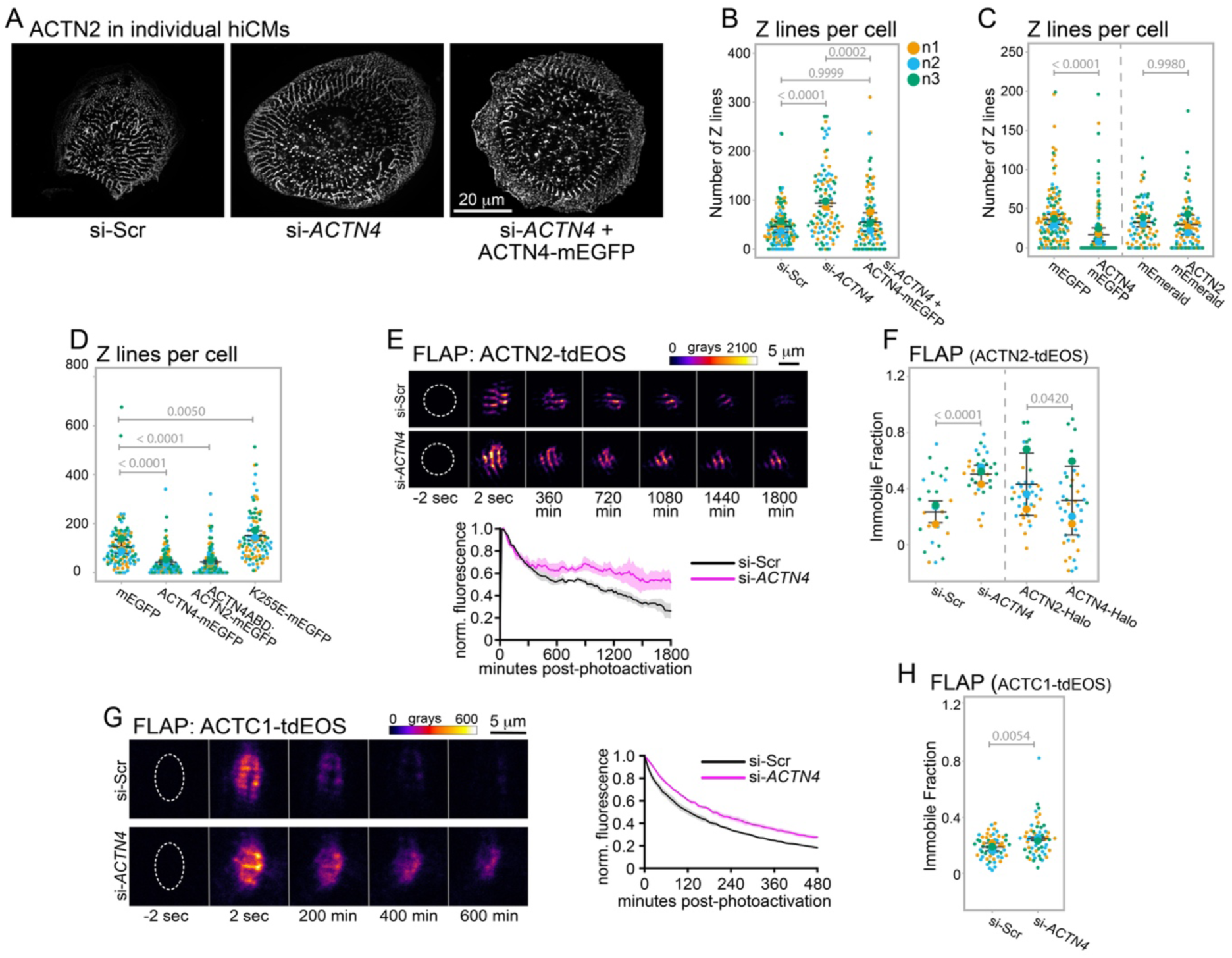
ACTN4 regulates sarcomere stability. **2A:** ACTN2 in hiCMs treated with non-targeting siRNA (si-Scr), *ACTN4*-targeting siRNA (si-*ACTN4*), or *ACTN4-*targeting siRNA + exogenously expressed ACTN4-mEGFP (si-*ACTN4* + ACTN4-mEGFP). **2B:** Z-lines per cell across the treatment conditions from 2A. **2C:** Z-lines per cell in hiCMs expressing mEGFP, ACTN4-mEGFP, mEmerald, and ACTN2-mEmerald. **2D:** Z-lines per cell expressing either mEGFP, ACTN4-mEGFP, ACTN4ABD:ACTN2-mEGFP, or K255E-mEGFP. **2E:** Top: representative montages of ACTN2-tdEOS fluorescence loss after photoconversion (FLAP) in a hiCM treated with either non-targeting (si-Scr) or *ACTN4-*targeting (si-*ACTN4*) siRNA. Bottom: population average curves of photoconverted fluorescence signal (normalized to initial photoconverted signal). Shaded regions represent standard error. **2F:** Immobile fractions of ACTN2-tdEOS calculated from individual curves of photoconverted fluorescence signal in hiCMs treated with non-targeting siRNA (si-Scr), *ACTN4-*targeting siRNA (si-*ACTN4*), or co-expressing either ACTN2-halo or ACTN4-halo. **2G:** Right: representative montage of ACTC1-tdEOS fluorescence loss after photoconversion (FLAP) in a hiCM treated with either non-targeting (si-Scr) or *ACTN4-*targeting (si-*ACTN4*) siRNA. Bottom: population average curves of photoconverted fluorescence signal (normalized to initial photoconverted signal). Shaded regions represent standard error. C) ACTC1 immobile fraction calculated within individual hiCMs treated with either non-targeting (si-Scr) or *ACTN4-*targeting (si-*ACTN4*) siRNA. **2H:** Immobile fractions of ACTC1-tdEOS calculated from individual curves of photoconverted fluorescence signal in hiCMs treated with non-targeting siRNA (si-Scr) or *ACTN4-*targeting siRNA.

Since experiments in non-muscle cells have found that ACTN4 destabilizes contractile actin stress fibers^41,42^, we hypothesized it may serve an analogous function in muscle and destabilize sarcomeres. Using a technique called fluorescence loss after photo-conversion (FLAP) we converted a sub-population of ACTN2-tdEOS (a photo-convertible fluorescent protein) to emit red light (instead of green) at the Z-disc and measured loss of red fluorescence intensity over time. FLAP revealed that a larger fraction of photo-converted ACTN2-tdEOS stayed bound at the Z-disc in ACTN4-depleted cells **(Fig. 2E-F)**. Conversely, overexpression of ACTN4-Halo, but not ACTN2-Halo, was associated with a decrease in the fraction of ACTN2-tdEOS which stayed bound at the Z-disc **(Fig. 2F)**. Because our data suggested ACTN4 regulates sarcomeres through actin cross-linking, we asked if ACTN4 depletion also impacted the stability of sarcomeric actin filaments. Similar to ACTN2-tdEOS, a larger fraction of photo-converted ACTC1-tdEOS (cardiac alpha-actin) stayed bound at the sarcomeres of ACTN4-depleted cells **(Fig. 2G-H)**. Taken together, these data show ACTN4 regulates turnover of actin filament cross-links at the CM Z-disc.

A recent computational model of non-muscle stress fibers predicted that reducing turnover of actin cross-linkers increases stress fiber contractility in non-muscle cells^43^. Therefore, we used traction force microscopy to ask if ACTN4 depletion – which reduced ACTN2 turnover – increased sarcomere contractility in CMs. We plated either ACTN4-depleted or control hiCMs on polyacrylamide substrates of physiological stiffness and measured the forces they generated during both contraction and relaxation. Forces generated on the substrate by ACTN4-depleted hiCMs were elevated while fully contracted but similar to controls while fully relaxed **(Fig. 3A-B)**. To distinguish if ACTN4 depletion increased contractility by increasing the number of sarcomeres or by increasing contractility of individual sarcomeres, we compared forces generated by cells of similar size, which our data suggested would have a similar number of sarcomeres **(Fig. S2G)**. Similarly-sized ACTN4-depleted hiCMs generated more force when fully contracted than control hiCMs **(Fig. 3C),** suggesting ACTN4 depletion increases contractility at the sarcomere-level.

**Figure 3:**
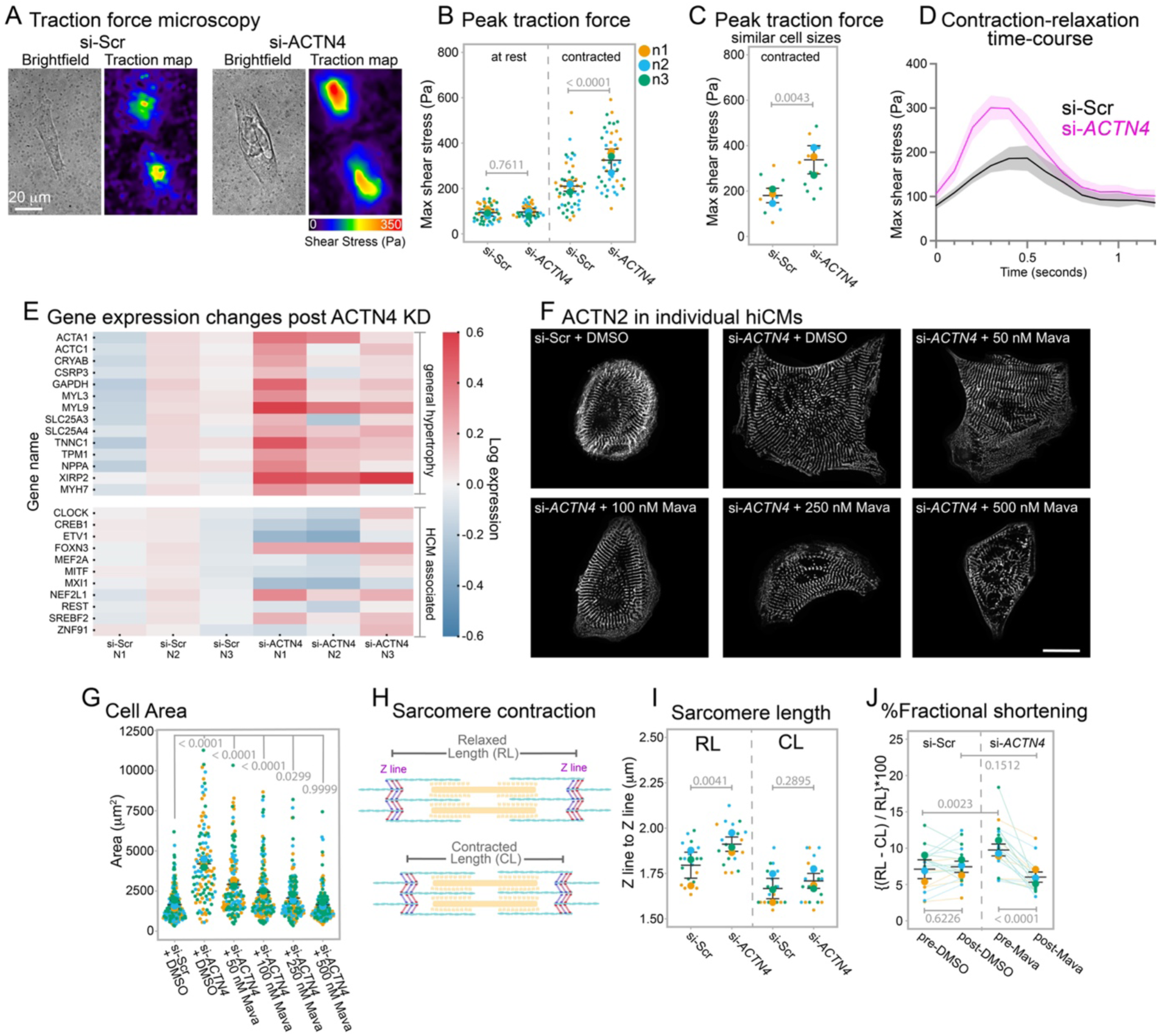
ACTN4 regulates sarcomere contractility. **3A:** Brightfield images and traction maps of a representative hiCM following treatment with either non-targeting (si-Scr) or *ACTN4-*targeting (si-*ACTN4*) siRNA. **3B:** Maximum sheer stress exerted by hiCMs while relaxed (passive force) or contracted (contractile force) treated with si-Scr or si*-ACTN4*. **3C:** Maximum contractile shear stress in hiCMs of similar areas. Areas were calculated from brightfield images. **3D:** Averaged maximum shear stress across one full contraction/relaxation cycle for hiCMs treated with si-Scr or si-*ACTN4*. **3E:** RNASeq of hiCMs following treatment with either non-targeting (si-Scr) or *ACTN4-*targeting (si-*ACTN4*) siRNA. Expression levels of general hypertrophy-and HCM-associated genes are normalized to si-Scr levels. **3F:** Representative examples of ACTN2 stains in hiCMs from each treatment group shown. **3G:** Cell area of hiCMs treated with either si-Scr+DMSO, si-*ACTN4*+DMSO, si-*ACTN4*+50nM mavacamten, si-*ACTN4*+100nM mavacamten, si-*ACTN4*+250nM mavacamten, or si-*ACTN4*+500nM mavacamten. Each small dot in each graph represents a single hiCM. **3H:** Schematic illustrating how sarcomere contraction impacts relaxed length (RL) and contracted length (CL), measured as the distance between Z-lines. **3I:** Sarcomere relaxed length (RL) and contracted length (CL) measured in live, beating hiCMs following treatment with non-targeting siRNA (si-Scr) or *ACTN4-*targeting (si-*ACTN4*) siRNA. **3J: %**Fractional shortening of individual sarcomeres in si-Scr (control) hiCMs pre-and post-DMSO vs. si-*ACTN4* (ACTN4-depleted) hiCMs following treatment with pre-and post-mavacamten (250nM).

Sarcomere mutations that increase contractility at the sarcomere-level cause hypertrophic cardiomyopathy (HCM)^12,44^. HCM mutations that increase force characteristically alter CM force curves by increasing maximum force and delaying/lengthening the return to baseline (i.e., relaxation)^45–48^. Therefore, we measured traction forces of control (si-Scr) and ACTN4-depleted (si-*ACTN4*) hiCMs during the entire contraction-relaxation cycle and aligned force curves to contraction onset. Force generation by ACTN4-depleted hiCMs was elevated during the contraction period but the return to baseline was unchanged **(Fig. 3D)**, suggesting ACTN4 depletion increased contractility through mechanisms distinct from HCM. To further test if ACTN4 depletion could be associated with HCM, we sequenced RNA isolated from hiCMs and compared the impact of ACTN4 depletion on the expression of genes that are either i) generally associated with hypertrophy or ii) specifically implicated in HCM. ACTN4 depletion resulted in broad upregulation of the general hypertrophy-associated genes^49–52^ but most of the specifically HCM-associated genes^52^ were downregulated **(Fig. 3E)**.

To determine if ACTN4 depletion-associated hypertrophy was a downstream consequence of increased sarcomere contractility, we asked if we could prevent hypertrophy by attenuating contractility during ACTN4 depletion. Reducing contractility with the MYH7 (β-myosin II) inhibitor, mavacamten^53^, attenuated the ACTN4 depletion-associated increase in cell size at low doses and completely prevented it at medium to high doses **(Fig. 3F-G)**. These data suggest ACTN4 depletion stimulates cellular hypertrophy by increasing sarcomere contractility. To directly test the hypothesis that ACTN4 depletion increases sarcomere contractility, we measured %fractional shortening, which quantifies the degree that individual sarcomeres shorten during contraction by computing percentage difference between contracted length (CL) and relaxed length (RL) **(Fig. 3H)**. Sarcomeres in ACTN4-depleted hiCMs had longer relaxed lengths than controls but similar contracted lengths **(Fig. 3I)**, suggesting ACTN4 depletion increases %fractional shortening of individual sarcomeres **(Fig. 3J)**. Interestingly, when we treated with the minimum mavacamten dose that fully prevented cellular hypertrophy (250 nM), it restored the elevated %fractional shortening of ACTN4-depleted sarcomeres to control levels **(Fig. 3J)**. Taken together, these data suggest ACTN4 depletion from CMs results in contractility-dependent cellular hypertrophy accompanied by increases in both sarcomere length and %fractional shortening.

### ACTN4 depletion *in vivo* results in contractility-dependent remodeling

Contractile force generation generally increases in cardiac muscle as resting cardiac sarcomere lengths increase between a working range of 1.8 μm to 2.3 μm^54–56^. Because ACTN4 depletion *in vitro* increased hiCM sarcomere length within this range we tested the hypothesis that ACTN4 depletion *in vivo* would increase cardiac contractility. The model organism zebrafish rapidly develops a two-chambered heart with both a nascent ventricle and atrium by ∼48 hours post fertilization^57^ (hpf). To first determine if ACTN4 was a sarcomere protein in the zebrafish embryo, we anesthetized 72 hpf embryos then fixed and stained ACTN4. Whole mount immunofluorescence revealed ACTN4 striations within the sarcomeres of the embryonic ventricle with little to no localization to the atrium **(Fig. 4A)**. Next, we depleted ACTN4 and measured relaxed sarcomere length in fixed ventricles at high magnification **(Fig. 4B, S3A-D)**. Sarcomere length in ACTN4-depleted embryos averaged 2.09 μm compared to 1.91 μm in control embryos **(Fig. 4B)**. These results matched our *in vitro* data but could also be explained by increased filling of the ventricular chamber by the atrium. Therefore, we depleted ACTN4 using two independent morpholinos and measured chamber size at low magnification. ACTN4 depletion did not change ventricle area but was associated with an increase in atrium area **(Fig. 4C-E)**.

**Figure 4:**
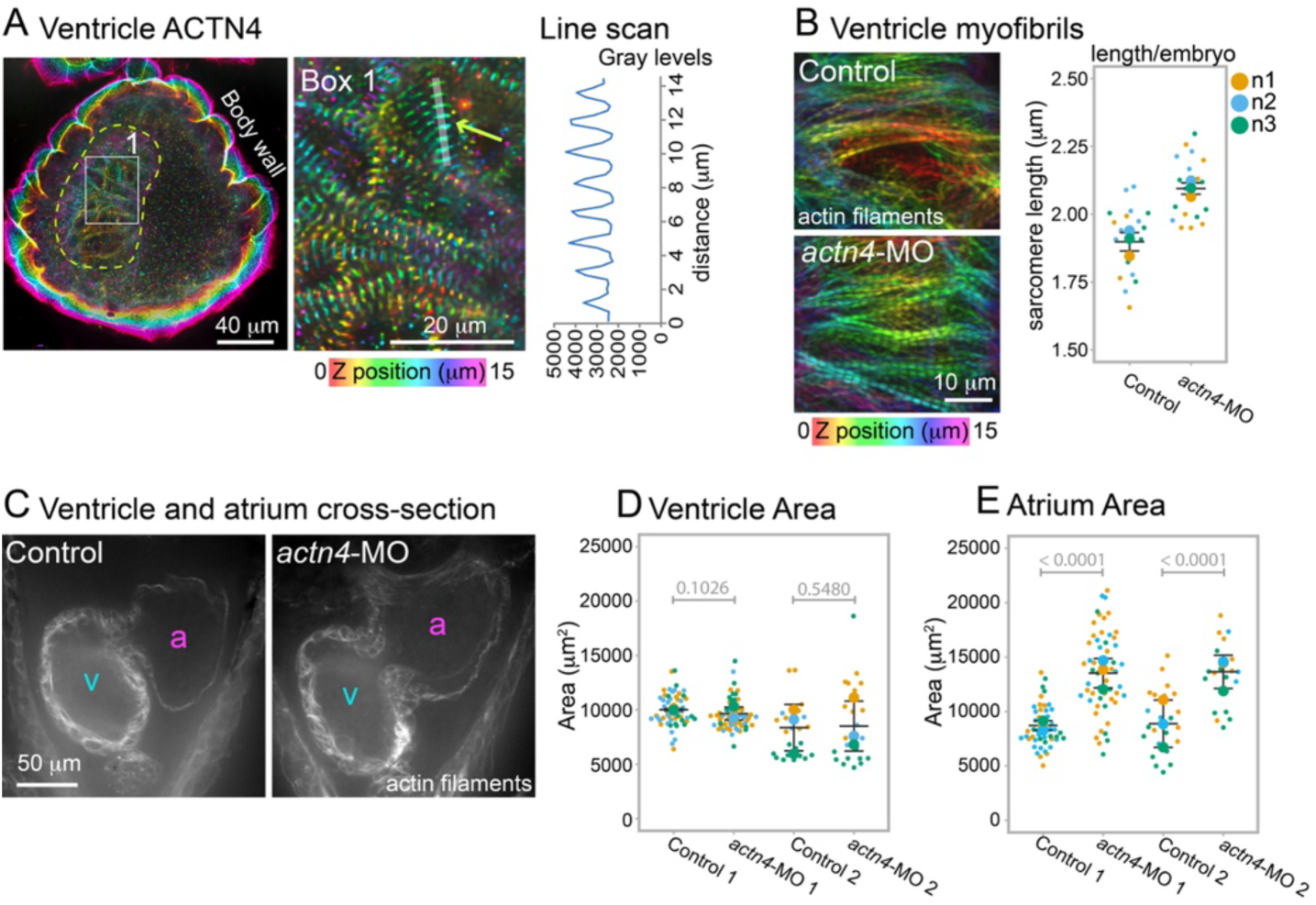
ACTN4 depletion *in vivo* increases sarcomere length and results in atrium enlargement. **4A:** Whole mount immunofluorescence stain of a zebrafish embryo showing actn4 striations visible in the ventricle at 72hpf. Box 1 shows high mag inlay. Line scans reveal ∼1.8um distance between striations, consistent with Z-line spacing. **4B:** Left: representative phalloidin stains of ventricle actin filaments in 72hpf zebrafish embryos that were either non-injected (control) or injected at one-cell stage with *actn4-* targeting morpholinos. Color denotes relative Z positions. Right: average relaxed sarcomere length of each zebrafish ventricle. **4C:** Cross-section of the myocardium of control and *actn4-*MO embryos. v and a denote ventricle and atrium, respectively. **4D:** Area of the ventricle of control (non-injected) or *actn4-*MO injected embryos **4E:** Area of the atrium of control (non-injected) or *actn4-*MO-injected embryos.

Atrium enlargement has been reported in zebrafish cardiomyopathy models, but is typically accompanied by pericardial edema with/without increased ventricle size and reduced ventricle contractility^58,59^. Unlike these models, ACTN4-depleted embryos had no overt organism-level phenotype nor edema **(Fig. S3E)** yet had elevated sarcomere lengths, suggesting the hypothesis that ACTN4 depletion increased ventricular contractility. To test this hypothesis, we mounted live embryos ventral-side down in slotted, stamp-cast agarose gels^60^ and assessed ventricular contractility by the classical measurement ejection fraction (EF), which represents the fraction of blood ejected from the ventricle during each contraction. EF in zebrafish can be estimated from chamber area curves of the full myocardial contraction-relaxation cycle **(Fig. 5A-B)** where EF values are generated from the areas measured when each chamber is fully relaxed (i.e., curve maximum) vs. fully contracted (i.e., curve minimum) **(Fig. 5B arrow + arrowheads)**. For each embryo we calculated EF at 72 hpf from maximum and minimum area of two full contraction-relaxation curves. Representative curves for each treatment group are shown aligned to the onset of ventricular contraction in **Fig. 5C** with representative images in **Fig. 5D**. ACTN4-depleted (*actn4-*MO + mApple) embryos had increased ventricular EF compared to non-injected (control) embryos **(Fig. 5E)** whereas atrial EF was unchanged **(Fig. 5F)**. Consistent with measurements in fixed embryos, ventricle area (measured at the fully relaxed state) was unchanged whereas atrium area was increased **(Fig. 5G-H)**. Genetic replacement with human ACTN4 (*actn4-*MO + hACTN4-mApple) concomitantly restored both ventricular EF and atrium area to control levels **(Fig. 5C-H)**, suggesting both phenotypes were a direct consequence of reducing ACTN4 levels. Furthermore, ventricle EF plotted directly against the area of the associated atrium revealed a positive correlation between EF and atrium area **(Fig. 5I; Pearson r=0.51)**, suggesting a positive association between ventricular contractility and atrium size.

**Figure 5:**
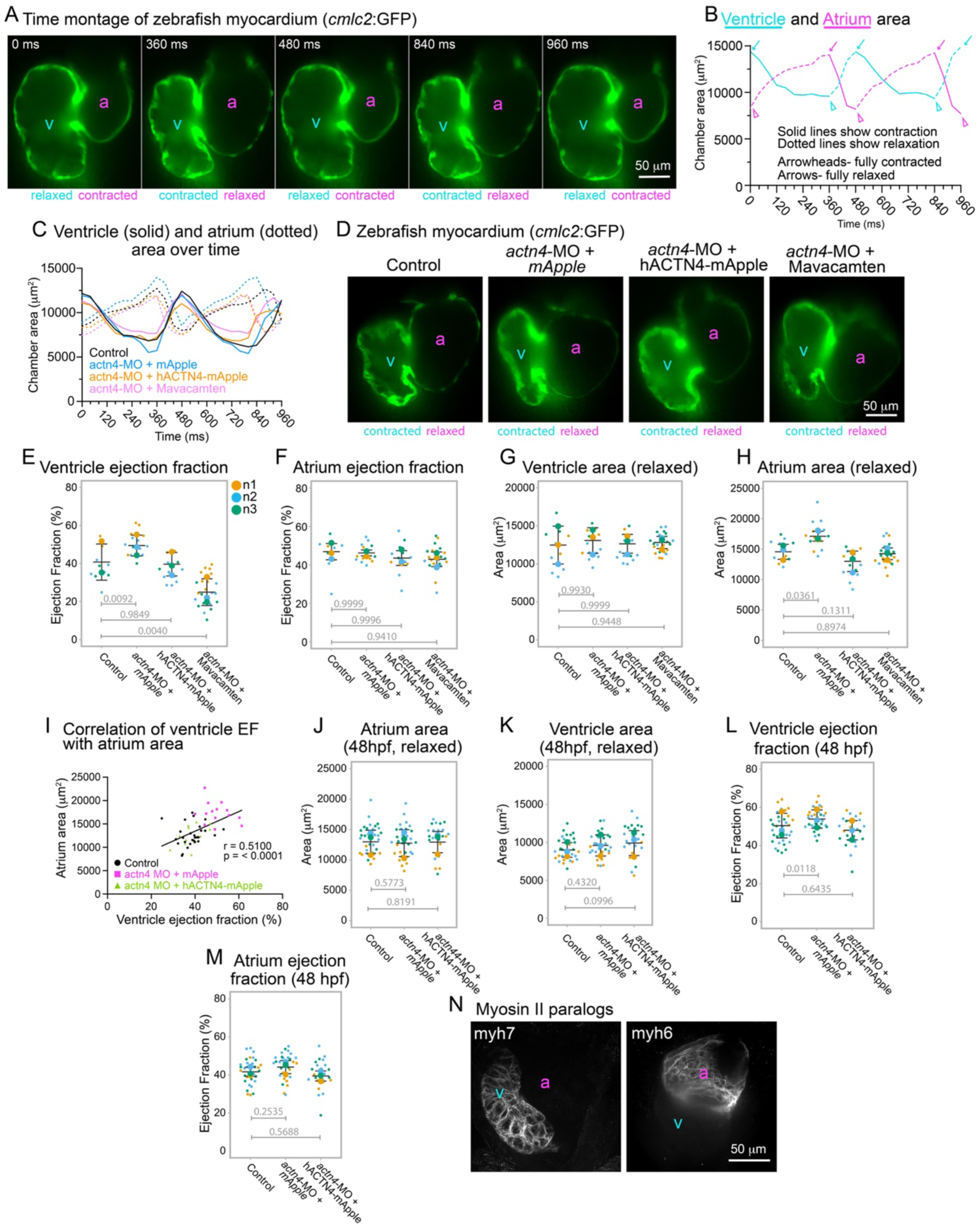
Ventricular contractility drives atrium enlargement following ACTN4 depletion *in vivo*. **5A:** Time montage of embryonic myocardium showing contraction-relaxation of a control (non-injected) heart. **5B:** Chamber areas across two full contraction-relaxation cycles for the embryo in (5A). **5C:** Chamber areas for control, *actn4-*MO + mApple, *actn4-*MO + hACTN4-mApple, and *actn4-*MO + 50uM mavacamten-treated (from 48hpf to 72hpf) embryos across two full contraction-relaxation cycles. **5D:** Representative single cross-sections showing fully contracted zebrafish embryonic ventricle (v) and fully relaxed atrium (a) at 72hpf. **5E:** Ventricle ejection fraction (EF) for all embryos within each treatment group from (5C). **5F:** Atrium ejection fraction (EF) for all embryos within each treatment group from (5C). **5G:** Ventricle area while relaxed for all embryos within each treatment group from (5C). **5H:** Atrium area while relaxed for all embryos within each treatment group from (5C). **5I:** Ventricle EF plotted against atrium area. r denotes Pearson’s correlation and solid line denotes linear regression. **5J:** Atrium area when fully relaxed at 48hpf, measured from inner chamber wall of live embryos either not injected (control) or injected with mix containing *actn4-MO + mApple* mRNA, or *actn4-MO + hACTN4-mApple* mRNA. **5K:** Ventricle area when fully relaxed at 48hpf from same treatment groups as (5J). **5L:** Ventricle ejection fraction at 48hpf from same treatment groups as (5J). **5M:** Atrium ejection fraction at 48hpf from same treatment groups as (5J). **5N:** Maximum intensity projections of zebrafish embryonic ventricle (v) and atrium (a) following whole mount IF staining at 72hpf.

We next asked if we could establish a causal relationship between increased ventricle EF and atrium size. Starling’s law of the heart states that acute stretch of the myocardial wall stimulates an acute increase in myocardial contractility^61,62^. Starling’s law inspired our first hypothesis, which was that the atrium enlarged first and ventricular contractility had increased acutely as a result of increased atrium-dependent filling. To test this hypothesis, we asked if the onset of atrium enlargement preceded the onset of increased ventricular EF in ACTN4-depleted embryos. Surprisingly, at 48 hpf, ACTN4-depleted embryos had no difference in atrial area **(Fig. 5J).** Ventricle area was also unchanged, suggesting ventricle filling was unchanged **(Fig. 5K)**. However, ventricular EF of ACTN4-depleted embryos was elevated at 48hpf unless we genetically replaced with hACTN4, with no change in atrium EF **(Fig. 5L-M)**. These data indicated that ventricular contractility increased independently of (and prior to) atrium enlargement, suggesting the alternative hypothesis that ventricular contractility contributed to atrium enlargement.

It was previously observed that mRNA encoding an ortholog of human β myosin II (MYH7; zebrafish myh7) is expressed exclusively in the developing ventricle of zebrafish embryos while an ortholog of α myosin (MYH6; zebrafish myh6) is expressed exclusively in the atrium^58,63^. First, we confirmed this through whole mount IF of 72 hpf zebrafish embryos using antibodies against human MYH7 or MYH6. Antibodies against human MYH7 exclusively stained the 72 hpf zebrafish ventricle while antibodies against human MYH7 exclusively stained the atrium **(Fig. 5N)**. We then asked if attenuating zebrafish myh7-dependent (i.e., ventricular) contractility starting at 48 hpf using the human β myosin II (MYH7) inhibitor, mavacamten, could prevent atrium enlargement in ACTN4-depleted embryos. Mavacamten treatment between 48hpf and 72hpf reduced EF specifically in the ventricle **(Fig. 5C-F)** and prevented the ACTN4-depletion associated increase in atrium size **(Fig. 5C-D, 5H)**.

Next, we asked if increasing zebrafish myh7-dependent (i.e., ventricular) contractility starting at 48 hpf using omecamtiv mecarbil (OM), which binds β myosin II similarly to mavacamten but increases β myosin II-mediated contractility^64,65^, would be sufficient to drive atrium enlargement in non-injected embryos by 72 hpf. Representative curves for control-DMSO and OM-treated embryos are shown aligned to the onset of ventricular contraction in **Fig. 6A** with representative images in **Fig. 6B**. OM increased EF specifically in the ventricle without changing ventricle size **(Fig. 6C-E).** Moreover, continuous exposure to OM starting at 48 hpf was sufficient to drive atrium enlargement by 72 hpf **(Fig. 6F)**, re-capitulating both temporally and structurally the phenotypic changes in the myocardium seen in ACTN4-depleted embryos. Taken together, we conclude that ACTN4 depletion in zebrafish embryos increases ventricular EF by 48 hpf which drives secondary atrium enlargement from 48 hpf to 72 hpf. Because atrium enlargement is not accompanied by an acute increase in atrial contractility^66,67^, which we would expect if the increase in size was related only to acute wall stretch, we conclude the change in atrium size reflects remodeling.

**Figure 6:**
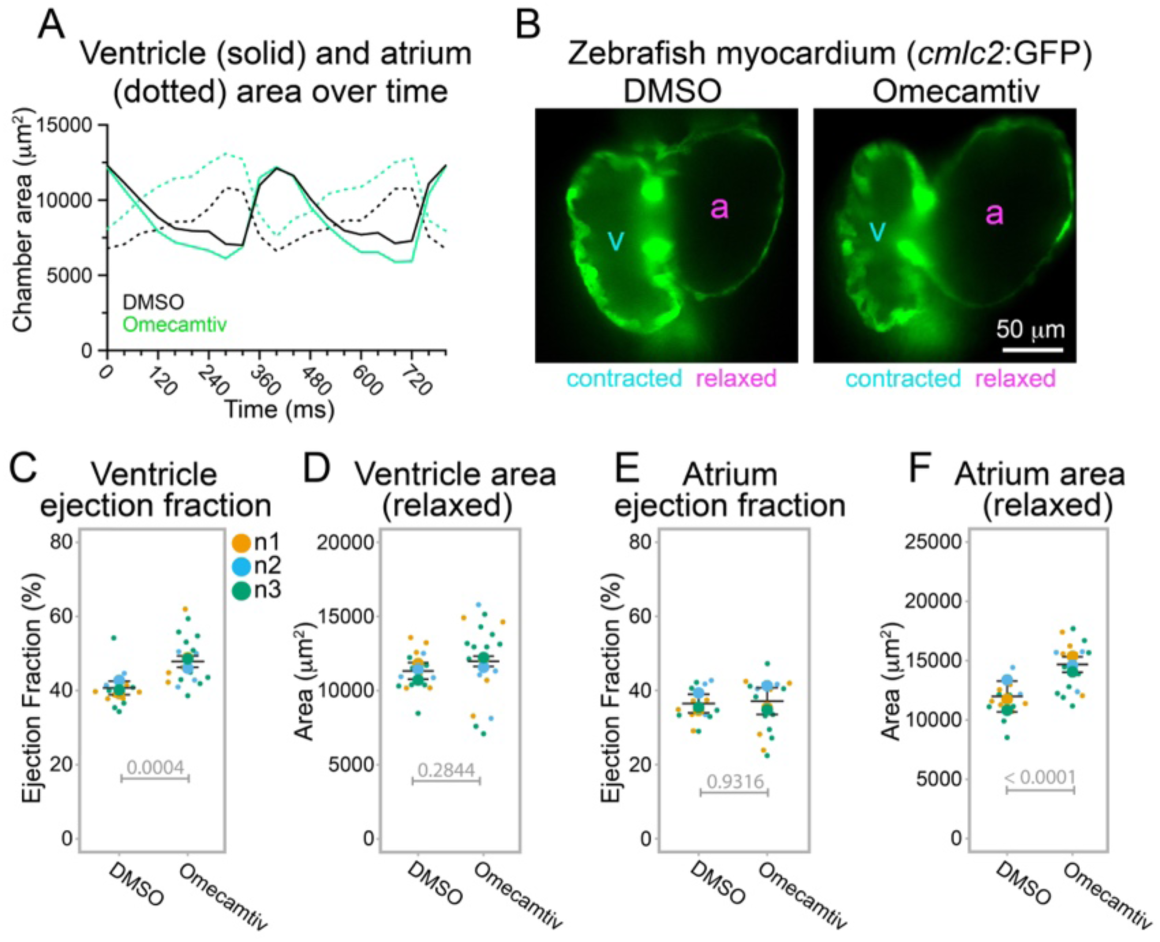
Increasing ventricular contractility is sufficient to drive atrium enlargement. **6A:** Chamber areas across two full contraction-relaxation cycles in embryos treated with DMSO control or 1uM omecamtiv mecarbil from 48hpf to 72hpf. **6B:** Representative single cross-sections showing fully contracted zebrafish embryonic ventricle (v) and fully relaxed atrium (a) at 72hpf following treatment with either DMSO or 1uM omecamtiv mecarbil from 48hpf to 72hpf. **6C:** Ventricle EF for all embryos within each treatment group from (6A). **6D:** Ventricle area within each treatment group from (6A). **6E:** Atrium EF within each treatment group from (6A). **6F:** Atrium area for all embryos within each treatment group from (6A).

To examine whether ACTN4 is linked with cardiac remodeling in humans we deployed computational genetics approaches. Given that ACTN4 depletion increased contractility, we hypothesized ACTN4 may have an association with heart failure with preserved ejection fraction (HFpEF), the end-stage heart condition associated with hypertrophic cardiomyopathy^14^. Using BioVU, Vanderbilt’s well-established biobank of de-identified electronic health records (EHRs)^68^, we tested 14 independent single nucleotide polymorphisms (SNPs) in ACTN4 for association with case status among 3,620 BioVU subjects with HFpEF (>57,000 controls). The strongest association with HFpEF was seen with the ACTN4 SNP where the G allele (MAF=29%) was associated with decreased HFpEF risk (OR 0.93; FDR 0.09). This variant, located within ACTN4 intron 1, is additively associated with reduced ACTN4 expression and has additionally been associated with altered splicing of ACTN4 in cardiac myocytes of the left ventricle (acquired through access to GTEx on 04/09/2024; https://ncbi.nlm.nih.gov/projects/gap/cgi-bin/study.cgi?study_id=phs000424.v9.p2). These data suggest that functional genetic variation in ACTN4 is associated with reduced HFpEF risk in humans.

## Discussion

A traditional approach in medical research is to identify a disease, then work backwards to understand its pathophysiology. As there is currently no known causative link between ACTN4 and any human cardiovascular disease, this traditional disease-oriented approach has failed to detect ACTN4 as a factor affecting heart biology. Our data nonetheless suggest ACTN4 regulates the core function of the heart and influences clinical outcomes related to heart failure. ACTN4 may have been overlooked in muscle for a multitude of reasons, including i) failure of classic studies of the ‘90s to detect any widely expressed actinins in muscle^69^, (ii) the abundance of ACTN2 in muscle in relation to the other paralogs^26^, (iii) long-term use of non-specific probes that likely recognize multiple actinin paralogs and, perhaps due to (i), (ii), and (iii), even (iv) a widespread assumption that ACTN2 is the only important actinin in muscle.

Our results suggest human cardiac myocytes contain ACTN4:ACTN2 heterodimers. The stoichiometric ratios of actinin homo-vs. putative heterodimers may carry functional significance in the cell and enable it to achieve different ends. Mechanistically, heterodimerization may provide a general means for a lower-expressed paralog (ACTN4) to more substantially impact the cell, by directly modulating the function of the higher-expressed paralog (ACTN2). In cardiac myocytes specifically, the impact of a small amount of ACTN4 might be even more pronounced since each ACTN4 monomer at the sarcomere Z-disc would cross-link two actin filaments (one at each end), and therefore exert regulatory control over two adjacent sarcomeres. It has been shown that mechanical manipulation of even a single sarcomere within a myofibril impacts the mechanical stability and force generation along the entire myofibril^70^. Thus our results together with these studies point to a role for ACTN4 in intrinsically tuning myofibril stability.

Our results challenge a decades-old paradigm that “non-muscle” paralogs of muscle-specific proteins are generally excluded from muscle sarcomeres and only function within muscle cells to facilitate sarcomere assembly^23,24,26–28^. We find that ACTN4 localizes directly to the cardiac sarcomere Z-disc where it controls turnover of a subset of canonical sarcomere proteins. More broadly our other experiments suggest that interrupting ACTN4-driven turnover not only dramatically impacts sarcomere-generated contractile force but that the cell can sense this impact and couples it with a hypertrophic response. Therefore, we propose that ACTN4-driven component turnover is an intrinsic regulatory feature of the sarcomere. Turnover could impact contractility indirectly by creating instability within filaments or more directly by regulating filament exchange specifically within contractile filaments^71^.

Our data suggest cardiac myocyte sarcomeres are regulated by ACTN4 actin cross-linking. Cross-linking by ACTN4 is profoundly impacted by the human kidney disease-causing mutation K255E^40,72^. Recent studies have reported a correlation between K255E-associated kidney disease and cardiomyopathy, even in children^73–75^. Our data suggest K255E-associated cardiomyopathy, rather than being driven solely by kidney dysfunction, may be at least partly attributable to changes at the sarcomere. Due to disease rarity, we were unable to procure echocardiogram or other heart-phenotypic data on K255E. ACTN4 actin cross-linking can be also regulated post-translationally by tandem phosphorylation at two residues within an N-terminal disordered region of ACTN4^76,77^. This N-terminal region is intact in our ACTN4ABD:ACTN2 chimera which reduced sarcomeres when expressed in hiCMs. Finally, several mutations in domains mediating ACTN2 dimerization have been linked to cardiomyopathies^78,79^. If these mutations impact ACTN2:ACTN4 heterodimerization, our data suggest contractile dysregulation of CMs may be a downstream consequence.

We demonstrate that the zebrafish embryo is pharmacologically compatible with human drugs targeting MYH7. It has been well-established by others in the field that the heart chambers of the zebrafish embryo are structurally responsive to perturbations that affect contractility of the adjoining chamber^58,67,80^. The mechanisms underlying structural remodeling appear to be especially sensitive to changes in contractility during development^81–84^. We were able to specifically inhibit or activate zebrafish ventricular contractility with no effect on atrium contractility. This approach may provide a powerful experimental paradigm for future studies probing the interplay between cardiovascular hemodynamics, contractile biophysics, and heart chamber remodeling, especially for understanding congenital heart disease-related defects related to disrupted biophysics of chamber formation^85,86^. In such studies, our data suggest the atrium could provide a direct readout of changes in ventricular contractility and provide a rapidly tractable system for hypothesis-driven investigation of biophysical mechanisms underlying normal and aberrant chamber development.

While the focus of cardiac research often concerns mechanisms of disease we characterize here a protein that appears to modulate contractility through mechanisms that are phenotypically distinct from those seen in disease. For instance while contractile cardiomyopathies typically lengthen the phase of contraction and reduce or do not change ejection fraction^87^, ACTN4 depletion-associated hypercontractility actually increased ejection fraction without apparent changes in contraction kinetics. Underscoring this difference was our finding that certain ACTN4 variants may be cardioprotective in humans, implicating ACTN4 in a type of physiological remodeling. Nevertheless, it remains conceivable that ACTN4 could also play a role during pathological remodeling especially in those disease states which either heighten or depress cardiac contractility. Two recent reports of differentially expressed genes in either HCM or pediatric DCM underscore this possibility, which found opposite directional effects on *ACTN4* expression^30,88^. Thus, our study implicates ACTN4 in physiological remodeling, while others in the literature suggest it may also play context-dependent roles during different disease states that could either be contributory or compensatory.

## METHODS

### Tables of reagents

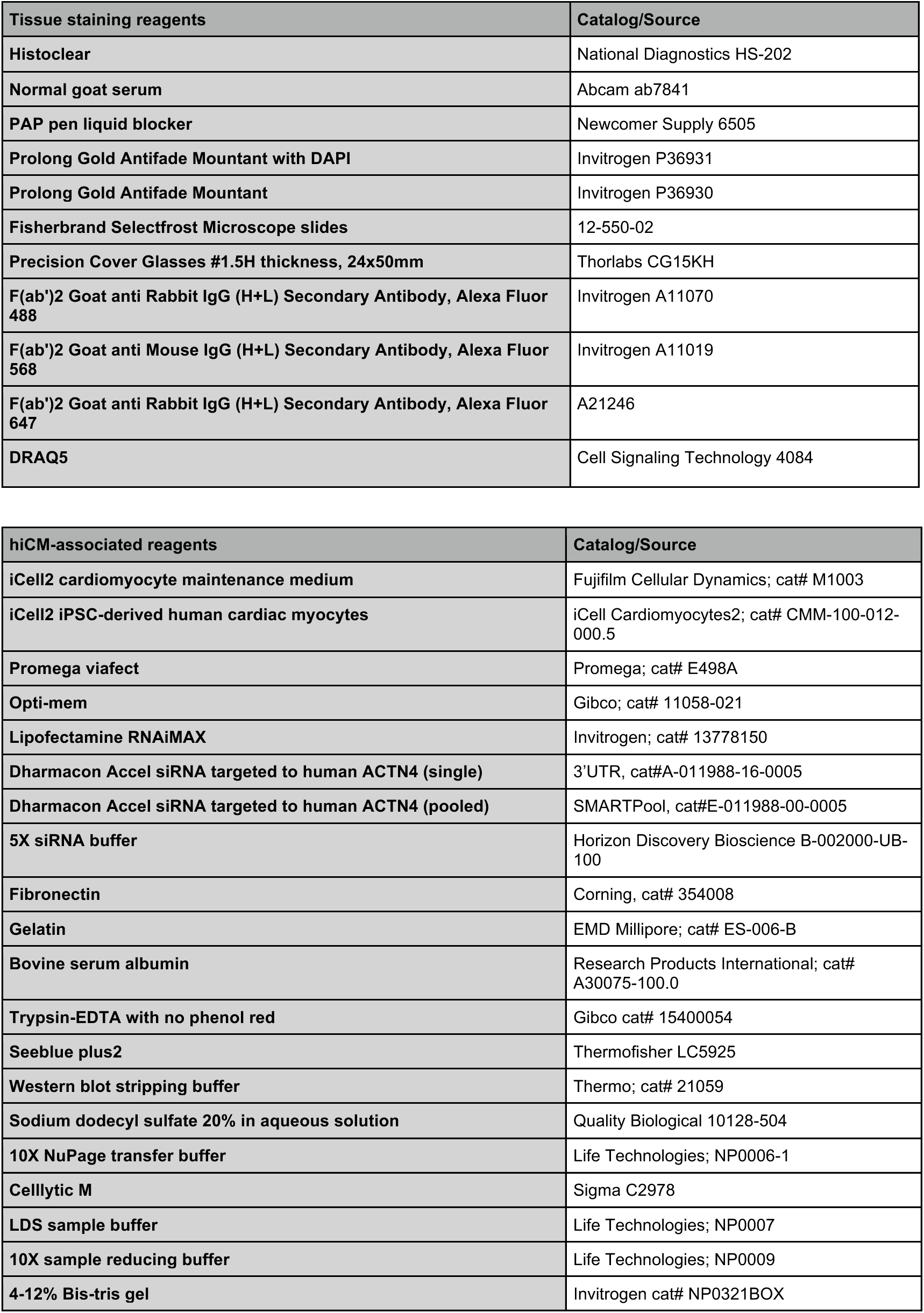
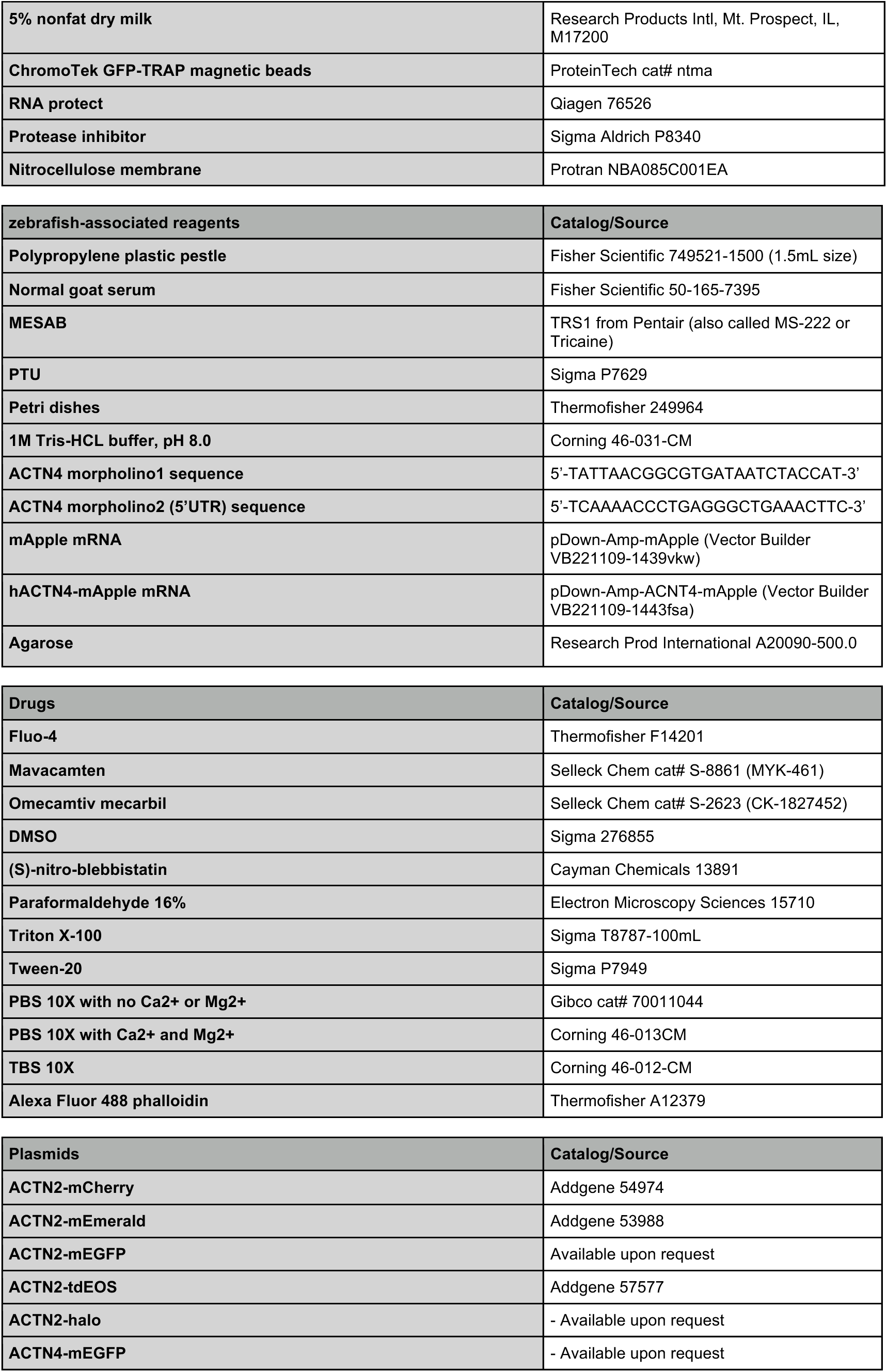
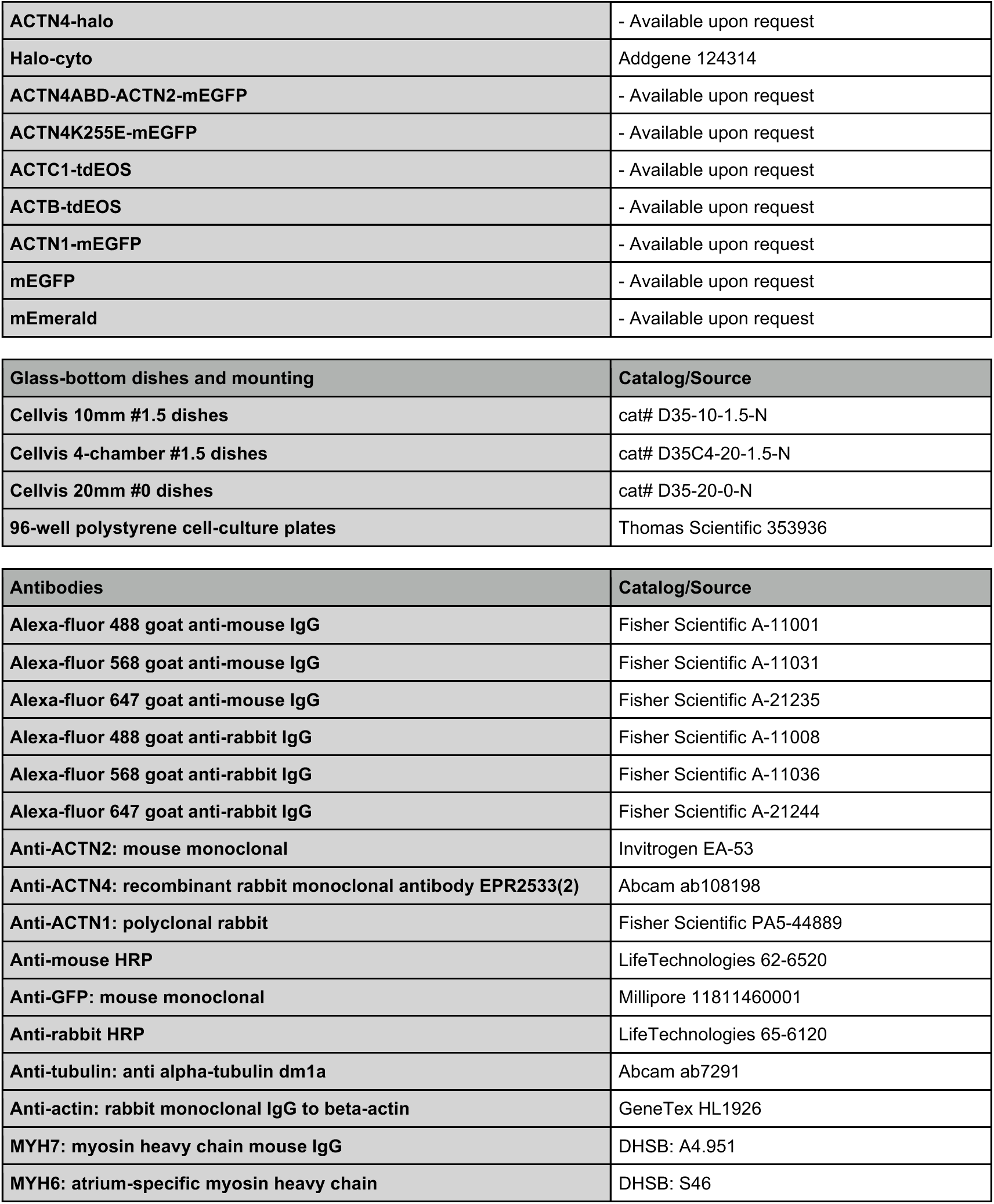

### Ethical approval

All animal studies were done in accordance with NIH, the US Department of Agriculture Animal Welfare Act, and the US Public Health Service Policy on Humane Care and Use of Laboratory Animals and were approved by Vanderbilt University Medical Center’s Institutional Animal Care and Use Committee. Zebrafish embryo experiments were conducted in accordance with M2100073-00-S2300172. Human tissue was acquired from the Vanderbilt Cardiology Core under the approval of IRB#240465.

### Tissue staining

Formalin-fixed paraffin-embedded slides were deparaffinized in Histoclear (National Diagnostics, HS-202) and rehydrated through a decreasing gradient of 100% ethanol in water. Antigen retrieval was performed using a rice cooker to boil samples for 1 hour in 10mM Tris, 0.5mM EGTA, pH 9 antigen retrieval buffer.

Slides were cooled to room temperature (RT), then washed 3x in phosphate buffered saline (1x PBS). A PAP pen was used to isolate tissue sections on the slide, and then slides were blocked for 1h at RT in 10% normal goat serum (NGS; ab7481 Abcam). Primary antibodies were diluted in 1% NGS at 4°C overnight. The next day slides were washed 3x in 1x PBS, then incubated in secondary antibody diluted in 1% NGS at RT in the dark. When appropriate, DRAQ5 was added to the secondary incubation to stain nuclei. Slides were then washed 3x in 1x PBS, then dehydrated in the reverse order of the ethanol gradient previously stated. Slides were mounted with Prolong Gold Antifade Mountant (P36930) with DAPI (P36931) when appropriate and sealed with clear nail polish.

### Cell culture

**<u>h</u>**uman **i**nduced pluripotent stem cell-derived **<u>c</u>**ardiac **<u>m</u>**yocytes (hiCMs) were purchased from Fujifilm Cellular Dynamics (iCell Cardiomyocytes^2^; cat# CMM-100-012-000.5) and cultured according to manufacturer instructions. Briefly, hiCMs were thawed into 96-well polystyrene cell-culture plates coated with gelatin (EMD Millipore; cat# ES-006-B) at a plating density of 50,000 cells/well. Cardiomyocyte maintenance media (Fujifilm Cellular Dynamics; cat# M1003) was exchanged 5 hours after thawing and subsequently replaced every 48 hours. Cells were maintained at 37 C and 5% CO2. hiCMs were then transferred to hiCM glass bottom 35 mm culture dishes for experiments using the following procedure: hiCMs in a well of a 96-well plate were washed twice with 100uL 1X PBS with no Ca^2+^ or Mg^2+^ (Gibco cat# 70011044), then trypsinized for 2.5 minutes in 40 μL 0.1% Trypsin-EDTA with no phenol red (Gibco cat# 15400054). Trypsin was quenched by the addition of 120 μL maintenance medium and re-suspended cells were pelleted via centrifugation at 0.2 rcf for 3 minutes. Following centrifugation, the supernatant was aspirated and the pellet was resuspended in maintenance medium. Re-suspended cells were pipetted onto 35 mm Cellvis (either 4-chamber; cat# D35C4-20-1.5-N or 10mm; cat# D35-10-1.5-N) cover glasses (or polyacrylamide gels; see: traction force microscopy section) coated with 10ug/mL human fibronectin (Corning, cat# 354008).

### siRNA and transfection

Treatments such as siRNA administration and transfections were performed on hiCMs in 96-well plates prior to transferring them to a 35 mm culture dish. siRNA treatments utilized Lipofectamine RNAiMAX (Invitrogen; cat# 13778-075); siRNAs used were Dharmacon Accel siRNA targeted to Human ACTN4 (3’UTR, cat#A-011988-16-0005; SMARTPool, cat#E-011988-00-0005) and non-targeting (cat#D-001910-01-05). siRNAs were briefly combined with RNAiMAX in opti-mem, then mixed into cardiomyocyte maintenance medium and added to hiCMs. Transfections utilized ViaFect (Promega; cat# E498A); ViaFect was briefly combined with 100ng plasmid DNA in Opti-Mem (Gibco; cat# 11058-021), then mixed into cardiomyocyte maintenance medium and added to hiCMs at a ratio of 0.6uL ViaFect:100ng DNA for 18-24 hours. Plasmids introduced in this study include an alpha-actinin 4ABD:alpha-actinin 2 chimera containing the N-terminal actin-binding domain (ABD) of alpha-actinin 4 (ACTN4 amino acids 1-266) in frame with the central rod and C-terminal EF-hand domains of alpha-actinin 2 (ACTN2 amino acids 255-894). Other plasmids used in the study were those encoding mEGFP, alpha-actinin 2-mCherry fusion, alpha-actinin 2-mEGFP fusion, alpha-actinin 4-mEGFP fusion, and alpha-actinin4K255E-mEGFP fusion.

### Immunofluorescence (IF)

24 hours after transferring hiCM to 35 mm culture dishes, cells were fixed in 4% PFA diluted into PBS for 20 minutes at room temperature, then permeabilized for 5 minutes in 4% PFA + 0.1% Triton X-100. For some antibodies, improvement in labeling quality occurred when 4% PFA fixation step was omitted in place of an immediate 45-minute permeabilization at room temperature. After permeabilization, hiCMs were washed 3 times in 5 mL PBS and stained with phalloidin for 2 hours at room temperature. Phalloidin was removed using 3 PBS washes of 5 mL each, then hiCMs were blocked in 5% bovine serum albumin (Research Products International; cat# A30075-100.0) in PBS (block buffer) at room temperature for 20 minutes. After blocking, hiCMs were labeled with primary antibodies diluted into block buffer at 4 C overnight. Primary antibody dilutions ranged from 1:100-1:200. The next morning, hiCMs were washed 3 times for 5 minutes each with block buffer, then labeled with secondary Alexa fluor-labeled antibodies diluted 1:100 into block buffer + DAPI at room temperature for 1 hour. After secondary labeling, hiCMs were washed 3 times in 5mL PBS and imaged in PBS.

### AlphaFold3 structure predictions

Protein structure predictions were generated with the AlphaFold3 server^89^ and visualized using the PyMOL molecular graphics system version 3.0 Schrodinger, LLC. All raw data output by AlphaFold3 is freely available at Mendeley Data doi: 10.17632/9xsdm9mvhx.1.

### Z-lines per cell and cell area analysis

hiCMs were allowed to spread for 24 hours then fixed with 4% paraformaldehyde and labeled with anti-alpha-actinin 2 (Z-lines) and DAPI (nucleus). DAPI staining was used to mark fields for imaging to avoid cherry-picking. Stained, fixed hiCMs were imaged at 60x magnification using an instant structured illumination (iSIM) microscope. Images were thresholded and binarized using yoU-Net and Z-lines per cell were quantified via the automatic quantification tool, sarcApp^90^. Cell area was computed manually using either actin (phalloidin) or alpha-actinin 2 labeling of the cell border. For experiments testing the effects of depletion or transient overexpression (or depletion, then transient overexpression) on Z-lines per cell, siRNA-or transfection-containing media was administered to hiCMs while they were in 96-well plates (i.e., prior to hiCM re-plating).

### Western blot

hiCMs in 96-well plates were briefly trypsinized in 0.1% trypsin-EDTA, then pelleted via centrifugation at 0.2rcf for 3 minutes. After aspirating the supernatant, pelleted hiCMs were resuspended in 400uL PBS and then re-pelleted via centrifugation at 0.3rcf for 4 minutes. The supernatants were aspirated and the remaining pellets were lysed in ice cold lysis buffer (CellLytic M [Sigma C2978] + protease inhibitor cocktail for 45 minutes-1hr. After lysis, samples were centrifuged at 13,000xRPM at 4C for 20 minutes and the supernatant was loaded into 4-12% bis-tris pre-cast gel lanes supplemented with LDS sample buffer (Life Technologies; NP0007) and sample reducing buffer (Life Technologies; NP0009) for electrophoresis at 100V. Gels were transferred to 0.45 micron nitrocellulose membranes (Protran NBA085C001EA) via wet membrane transfer at 100V for 1 hr 15 minutes in NuPage transfer buffer (Life Technologies; NP0006-1) supplemented with 10% methanol and 0.1% SDS. After transfer, membranes were blocked in 5% nonfat dry milk (Research Products International; cat# M17200-500.0) + Tris-buffered saline with Tween-20 (TBST, 10X TBS from Corning 46-012-CM; Tween-20 from Sigma P7949) for 1 hour, then incubated with primary antibodies diluted in 5% milk + TBST at 4C overnight. The next morning, membranes were washed 3 times 5 minutes each with TBST, then incubated with HRP-conjugated secondary antibodies at room temperature for 1 hr. Membranes were then washed 3 times 5 minutes with TBST and imaged via chemiluminescence.

### Validation of antibody specificity

All immunofluorescence (including whole-mount), co-immunoprecipitation, and western blot experiments utilized the anti-alpha-actinin 4 recombinant rabbit monoclonal antibody EPR2533(2) (Abcam; ab108198). To verify specificity of EPR2533(2) for alpha-actinin 4, experiments were conducted to test EPR2533(2) for potential cross reactivity to alpha-actinin 2. hiCMs were transfected with either i) a plasmid containing an alpha-actinin 2-mEmerald open reading frame or ii) a vehicle for 24 hours, then pelleted, lysed, and immunoblotted with an anti-alpha-actinin 2 primary (mouse monoclonal EA-53) and an HRP-conjugated goat anti-mouse secondary. After imaging, the blot was stripped (Thermo; cat# 21059) and re-blotted using an anti-alpha-actinin 4 (EPR2533[2]) primary and an HRP-conjugated goat anti-rabbit secondary.

### Co-immunoprecipitation

hiCMs were transfected with either i) ACTN4-mEGFP, ii) mEGFP, or iii) a vehicle for a beads-only control, then briefly trypsinized in 0.1% trypsin-EDTA and pelleted via centrifugation at 0.2rcf for 3 minutes. After aspirating the supernatant, pelleted hiCMs were resuspended in 400uL PBS and then re-pelleted via centrifugation at 0.3rcf for 4 minutes. The supernatants were aspirated and remaining pellets were lysed on ice for 15 minutes in NP-40 lysis buffer (6mM Na_2_HPO_4_, 4mM NaH_2_PO_4_, 1% NP-40, 150mM NaCl, 2mM EDTA, 50mM NaF, 5ug/mL leupeptin, 0.1mM Na_4_VO_4_) + protease inhibitors. 5uL ChromoTek GFP-TRAP magnetic beads (Proteintech cat# gtma) were added to cleared cell lysates, tumbled at 4 C for 1 hr, then washed 3 times with NP-40 buffer. Proteins were eluted from beads via boiling for 5 minutes in SDS sample buffer, then separated on 4-12% bis-tris gels (Invitrogen cat# NP0321BOX) and transferred to nitrocellulose membranes (Protran; cat# NBA085C001EA). Membranes were blocked with TBST + 5% nonfat dry milk then immunoblotted with anti-alpha-actinin 2 (Invitrogen EA-53; mouse monoclonal) and anti-actin (GeneTex HL1926; rabbit monoclonal) primaries followed by HRP-conjugated goat anti-mouse and anti-rabbit secondaries. After imaging via chemiluminescence, membranes were stripped (Thermo; cat# 21059) and re-blotted with anti-alpha-actinin 4 (Abcam EPR2533[2]; rabbit monoclonal) and anti-GFP (Millipore 11811460001; mouse monoclonal) primaries followed by HRP-conjugated goat anti-mouse and anti-rabbit secondaries for re-imaging.

### Fluorescence Loss After Photoactivation (FLAP)

To investigate if ACTN4 depletion stabilizes sarcomeres, myocytes were transfected on the final media change post-ACTN4 depletion. To investigate if transient ACTN4 overexpression destabilizes sarcomeres, previously untreated myocytes were co-transfected with two constructs simultaneously – one containing ACTN2 and another containing either an ACTN4-fusion protein or just the fusion protein alone (the fusion protein chosen being specific to the given experiment). For FRAP experiments post-depletion, transfection constructs contained either ACTN2-mEmerald (for measuring actinin2 turnover) or ACTC1-mGFP (for measuring cardiac actin turnover). For photoactivation experiments post-depletion, transfection constructs contained either ACTN2-tdeos or ACTC1-tdeos.

The day after transfection, myocytes were plated onto 4-quadrant Cellvis dishes coated with 10ug/mL human fibronectin and allowed to spread for either 24 hours (ACTN2) or 48 hours (ACTC1). Following selection of 20 fields for each treatment group, a 405-nm laser was used for photobleaching/photoactivation of a selected region of interest, followed by imaging at temporal intervals that were deemed appropriate for the given experiment (determined empirically).

### Traction force microscopy

To investigate if ACTN4 depletion increases sarcomere-dependent force output, ACTN4-depleted (and control) hiCMs were trypsinized and re-plated onto polyacrylamide gel substrates of stiffness 8.7 kPa to mimic the embryonic myocardium. Polyacrylamide gels for TFM were fabricated as described elsewhere^91^. Briefly, the ratios of acrylamide to bis-acrylamide in solution were 7.5% acrylamide (40% w/v solution, Bio-Rad, Hercules, CA) and 0.25% N,N′-methylene-bis-acrylamide (2% w/v solution, Bio-Rad) to tune gel stiffness to 8.6 kPa. PA gels were embedded with 0.5 μm diameter red fluorescent beads (ThermoFisher Scientific). Substrate surfaces were functionalized using N-6-((acryloyl)amido)hexanoic acid ^92^ and coated with 0.2 mg/ml fibronectin embedded with fluorescent 647-nm beads.

Traction force microscopy was performed as previously described^92^. Myocytes deform the gel substrate when beating, displacing the beads. Bead displacement during a full contraction cycle was captured for individual myocytes 24 hours post-plating using a spinning disk confocal 40X magnification objective. Myocytes were then completely removed from the gel using 0.5% trypsin-EDTA to capture bead position of the fully relaxed gel. Bead position of individual myocytes from relaxation-contraction-relaxation were assembled into image stacks that were fully interleaved with bead position of the relaxed gel. Fully interleaved image stacks were processed using the PIV and FTCC FIJI plugins (Qingzong Tseng) with an assumed Poisson ratio=0.5.

### hiCM mavacamten rescue experiments

To investigate if a reduction in contractility rescues ACTN4 depletion phenotypes in hiCMs, media containing ACTN4 siRNA was supplemented with the cardiac beta myosin II small molecule inhibitor, mavacamten (Selleck Chem cat# S-8861). Mavacamten was mixed into siRNA-containing media at either 50nM, 100nM, 250nM, or 500nM concentrations and was washed off at the conclusion of siRNA treatment. hiCMs were then re-plated using the standard protocol (see: Re-plating assay) and Z-lines and cell area were analyzed (see: Z-lines per cell and cell area analysis).

### hiCM fractional shortening experiments

To investigate how ACTN4 depletion impacts sarcomere contractile mechanics and by what mechanism mavacamten rescues any change in contractile mechanics, the Z-lines of ACTN4-depleted and control hiCMs were imaged live while beating using differential interference contrast (DIC) imaging before and after drug addition. First, pre-drug beating movies were acquired by selecting stage points of beating siScr and si-*ACTN4* hiCMs. Media was then carefully removed and replaced with new media supplemented with either DMSO (for siScr hiCMs) or 250nM mavacamten in DMSO (for si-*ACTN4* hiCMs). Following a 15-minute equilibration period, post-drug beating movies were acquired by imaging the same hiCMs in media containing the drug. Pre-and post-drug stacks were aligned such that the same sarcomeres could be measured before and after drug administration.

To analyze contractile mechanics, sarcomere length across time was calculated by measuring the distance between two Z-lines during relaxation-contraction-relaxation. Z-line distance was measured from kymographs of 8-pixel width line scans drawn across two Z-lines during beating. Distance was calculated by first measuring number of pixels between Z-line gray-level intensity peaks, then converting pixels to microns. Relaxed length was calculated as the average distance between Z-lines in microns across five relaxed timepoints. Contracted length was calculated as the minimum micron distance during the acquisition. Maximum length was calculated as the maximum micron distance during the acquisition. Fractional shortening was calculated using the equation %fs=(1-(contracted length/relaxed length))*100.

### Zebrafish morpholino injections

Zebrafish (either wildtype strain AB or containing the *cmlc2:*gfp transgene) were set up as male-female pairs into individual tanks set with clear dividers overnight. The next morning, dividers were removed and fertilized embryos were harvested and injected at the one-cell stage. Injections were performed with micro-needles cut from glass capillary tubes. Injection mixes contained a mixture of Danieau solution, morpholino, and mRNA encoding either mApple (injection control) or ACTN4-mApple (rescue). After injection, embryos were moved to petri dishes containing egg water + methylene blue (anti-fungal). Unfertilized embryos were discarded 4-8 hours post-fertilization. Viable embryos were moved to egg water containing PTU at 24 hours post-fertilization.

Morpholinos were designed and ordered from GeneTools. ACTN4 morpholinos were either i) translation-blocking (sequence: 5’-TATTAACGGCGTGATAATCTACCAT-3’) or ii) targeted to the ACTN4 5’UTR (sequence: 5’-TCAAAACCCTGAGGGCTGAAACTTC-3’). Injection mixes contained 4ng morpholino unless otherwise stated. Morpholino injections also contained 100ng/uL mRNA encoding either mApple (injection control) or human ACTN4-mApple (hACTN4-mApple; rescue). mRNA was amplified from ACTN4-mEGFP by PCR amplification and purification (28104, Qiagen; Hilden, Germany). The cDNA was inserted into pGEM-T using TA subcloning (A1360, Promega; Madison, WI). Following restriction enzyme digestion, the insert was moved into pCS2+ and linearized by NotI digestion. Synthesized capped mRNA transcripts were produced *via in vitro* transcription with mMESSAGE mMACHINE SP6 transcription kit (AM1340, ThermoFisher Scientific; Waltham, MA). The hACTN4-mApple mRNA contained the ACTN4 coding sequence matching the primary listed sequence for human ACTN4 on Ensemble (ACTN4-201 – ENST00000252699) with the C-terminal STOP codon mutated to a run-on sequence connected to a linker region plus the mApple coding sequence.

### Zmold preparation

The Zmold design file was kindly provided by Yijie Geng (University of Utah ^60^. The file was delivered as the filetype. step and converted to the filetype.stl using the STAMPA3D online conversion tool. The unedited.stl file was 3D-printed using tough resin at the Vanderbilt Institute of Nanoscale Science and Engineering. The resulting Zmold resin contained 8 “teeth”, each of which can hold a single zebrafish embryo up to 7 days post fertilization. Zmold resin was positioned in the center of a Cellvis 35mm glass-bottom dish with 20mm microwell #0 cover glass (catalog D35-20-0-N) and covered with 3mL of hot 2% agarose. Agarose was allowed to cool and solidify and Zmold resin was removed with tweezers carefully as not to introduce an air bubble between the solidified agarose and the cover glass. Zmold casts were then used to mount zebrafish for imaging.

### Zebrafish mounting

For fixed imaging, zebrafish were mounted in Zmold 2% agarose casts ventral side down (i.e., with hearts facing the cover glass). For live imaging, embryos from each treatment group were removed from 28°C incubator pipetted in 1mL suspensions into petri dishes containing 9mL of egg water supplemented with 750uM (200mg/mL) tricaine, resulting in an effective tricaine conc of 675uM (180mg/mL). After a 10 minute equilibration period, embryos were pipetted in 0.5mL suspensions into Zmold, resulting in an approximate final effective tricaine concentration of ∼100uM (25mg/L), and mounted for imaging. Embryos were imaged at room temperature (22-23°C) after an additional 10 minute equilibration period. All fish regardless of treatment group were anesthetized using the same Tricaine stock. Positioning was performed using mini-tweezers and aimed at stationing the beating heart as close to the cover slip as possible without damaging the embryo. Fish imaged at 48hpf were dechorionated prior to anesthetization and could be gently removed from Zmold after imaging using mini-tweezers and returned to the incubator to be re-mounted and re-imaged at 72hpf.

### Zebrafish drug treatments

For both mavacamten and omecamtiv mecarbil treatments, zebrafish embryos were transferred to egg water + PTU containing the drug diluted to the desired concentration in DMSO at 48 hpf. Control fish were transferred to egg water + PTU containing DMSO at an equivalent concentration. Just prior to mounting and imaging at 72 hpf, fish were transferred to egg water + PTU + drug containing 100 μM (25 mg/L) tricaine.

### Live cmlc2:gfp zebrafish imaging

Cellvis glass-bottom dishes containing zebrafish mounted in Zmold agarose casts were positioned on a Nikon spinning disk stage set up for live cell imaging. Imaging was performed using a 40X magnification oil immersion objective lens. Fish were imaged using both the brightfield and FITC channels. Brightfield imaging was Koehler-adjusted at a temporal sampling rate of 30 ms. FITC imaging was performed at a temporal sampling rate of 40 ms. For each heart, live beating movies were captured as follows: an acquisition was set up to continuous recording and a field of view was swept from the bottom of the coverslip through (when possible) the entire heart, pausing at the center of both chambers. If the boundaries of one or both chambers fell outside the working distance of the objective, imaging concluded at the furthest point from the objective that could be imaged without touching the coverslip.

### Chamber area curve generation and ejection fraction calculation

For each live, beating heart acquisition, the series of image slices containing the largest relaxed cross-section was identified for each chamber. Chamber area was quantified across two subsequent contraction-relaxation cycles using FIJI freehand selections of the chamber inner wall. Plots of chamber area across time were generated for beat rate-matched fish aligned to the onset of contraction for each chamber. Relaxed area (max area) and contracted area (min area) were derived from individual max and min values within each curve. Ejection fraction was calculated from the equation ((max area)-(min area)/max area)*100.

### Zebrafish phalloidin staining

A slightly modified version of a protocol demonstrated by Goody and Henry (http://bio-protocol.org/e786) was used for phalloidin staining. Zebrafish embryos (72 hpf) were fully anesthetized in Tricaine on ice, then fixed in 4% PFA in PBS rocking overnight at 4 C. The next day, embryos were washed and gently rocked 3 times for 5-10 minutes each in PBT (PBS + 0.1% Tween). After the last wash, embryos were permeabilized in PBS + 2% Triton X-100 for 3 hours rocking at room temperature. Permeabilized embryos were transferred to tubes containing 15uL phalloidin: 185uL PBS + 2% Triton X-100 and rocked overnight at 4°C. Tubes were kept in the dark for all subsequent steps. The next morning, embryos were washed 5 times for 5 minutes in PBT, then stained with DAPI and mounted for imaging.

### Zebrafish sarcomere length quantification

At 72hpf, non-injected and ACTN4 morpholino-injected zebrafish embryos were incubated in 10 μM blebbistatin (Cayman Chemicals 13891) at 28C for 15 minutes to relax the heart and stop contractions^93^, then fixed overnight in 4% PFA containing 10 μM blebbistatin. Following phalloidin staining + DAPI (see: Zebrafish phalloidin staining), embryos were mounted in Zmold 2% agarose casts (see: Zmold preparation and Zebrafish mounting) ventral side down and imaged at 60X magnification using a Nikon CSU-W1 SoRa. Sarcomere length was calculated as the average Z-line spacing per animal. Z-line spacing was calculated from line scans across phalloidin-stained, ventricular myofibrils, which revealed fluorescence peaks at Z-lines. Z-line spacing was measured as the pixel distance between peaks, converted to microns.

### Zebrafish whole-mount antibody staining

A slightly modified version of a protocol demonstrated by Inoue *et al.* ^94^ was used for whole mount staining. Zebrafish embryos (72 hpf) were fully anesthetized in Tricaine on ice, then fixed in 4% PFA in PBS rocking overnight at 4°C. The next day, embryos were washed and gently rocked 3 times for 5-10 minutes each in PBT (PBS + 0.1% Tween). Embryos were then incubated in 150 mM Tris-HCl at pH 9.0 for 5 minutes, then heated at 70°C for 15 minutes. After heating, embryos were washed twice in PBT for 5 minutes each, then twice in deionized water for 5 minutes each, followed by submersion in ice cold pure acetone at-20 C for twenty minutes. After cold treatment, embryos were again washed twice in deionized water for 5 minutes each, then twice in PBT for 5 minutes each. Embryos were blocked for three hours at room temperature in block solution containing 10% natural goat serum, 2% triton X-100 and 1% BSA in PBT. After blocking, embryos were rocked at 4°C for three days in I-buffer solution (same as block solution except containing only 1% natural goat) containing the desired primary antibodies. After 3-day primary incubation, embryos were washed three times for one hour each in solution containing 10% goat serum and 2% Triton X-100 in 1X PBS, then two times for one hour each in solution containing 2% Triton X-100 in 1X PBS. Embryos were then incubated in secondary antibody plus DAPI at 4C in the dark for 2.5 days at 4C. After 2.5-day incubation, Embryos were again washed three times for one hour each in solution containing 10% goat serum and 2% Triton X-100 in 1X PBS, then two times for one hour each in PBT. Primary antibody dilutions used were 1:100-1:200 and secondary antibody dilutions used were 1:50. DAPI was diluted to 1ug/mL.

### Zebrafish western blotting

Zebrafish embryos (72hpf) were fully anesthetized in Tricaine on ice, then 12-15 embryos were transferred to tubes containing ice cold lysis buffer (CellLytic M [Sigma C2978] + protease inhibitor cocktail. Embryos were crushed in lysis buffer using disposable pellet pestles, then shaken gently on ice for 15 minutes. After 15 minutes, embryos were centrifuged at 13,000xRPM at 4C for 20 minutes and the supernatant was loaded into 4-12% bis-tris pre-cast gel lanes supplemented with LDS sample buffer (Life Technologies; NP0007) and sample reducing buffer (Life Technologies; NP0009) for electrophoresis at 100V. Gels were transferred to 0.45 micron nitrocellulose membranes (Protran NBA085C001EA) via wet membrane transfer at 100V for 1 hr 15 minutes in NuPage transfer buffer (Life Technologies; NP0006-1) supplemented with 10% methanol and 0.1% SDS. After transfer, membranes were blocked in 5% nonfat dry milk (Research Products International, Mt. Prospect, IL, M17200) + Tris-buffered saline with Tween-20 (TBST, 10X TBS from Corning 46-012-CM; Tween-20 from Sigma P7949) for 1 hour, then incubated with primary antibodies diluted in 5% milk + TBST at 4C overnight. The next morning, membranes were washed 3 times 5 minutes each with TBST, then incubated with HRP-conjugated secondary antibodies at room temperature for 1 hr. Membranes were then washed 3 times 5 minutes with TBST and imaged via chemiluminescence.

### Zmold for use of live cardiac contraction-relaxation imaging

We bypassed traditional methods of zebrafish imaging in this study in favor of the Zmold metho ^60^. Zmold was originally designed for zebrafish brain imaging but we show that it is quite versatile as a simple imaging solution for zebrafish as it was originally designed for 7-day old fish larvae positioned dorsally. Here, we show it can also be used to reliably image 3-day old embryos positioned ventrally. Zmold enabled us to perform what was essentially high-throughput imaging of both live and fixed zebrafish hearts using a spinning disk confocal, however the use of Zmold in this manner should be compatible with many other imaging modalities. We found the primary strength of Zmold to be the ability to rapidly mount samples in identical orientation – this greatly increased the number of animals we could feasibly image which facilitated quantitative measurements of phenotypes like wall thickness that may have gone undetected in smaller populations. We additionally note that Zmold was compatible with longitudinal measurements as we could image a live population at 2-days post-fertilization, then carefully extract and re-image the same population at 3-days post-fertilization. Zmold as it was used here may not be compatible for some applications due to the limited objective working distance which combined with light scattering prevented in-focus volumetric imaging of most zebrafish hearts. We envision that inverted imaging modalities which utilize a liquid immersion objective would be particularly strong in combination with Zmold.

### Human and mouse tissue stains

Formalin-fixed paraffin-embedded (FFPE) slides were initially deparaffinized using Histoclear (National Diagnostics, HS-202) followed by rehydration through a gradient of decreasing ethanol concentrations in water. For antigen retrieval, slides were boiled for 1 hour in a rice cooker containing 10 mM Tris and 0.5 mM EGTA buffer at pH 9. Once cooled to room temperature, slides were rinsed three times with phosphate-buffered saline (1X PBS). Tissue sections were outlined with a PAP pen and slides were blocked for 1 hour at room temperature using 10% normal goat serum (NGS; Abcam ab7481). Primary antibodies were diluted in 1% NGS and incubated overnight at 4°C. The following day, slides were washed three times with 1X PBS then incubated with secondary antibodies diluted in 1% NGS and kept at room temperature and in the dark. If needed, DRAQ5 was added during this step to stain the nuclei. After washing slides three times with 1X PBS, slides were dehydrated using a reverse ethanol gradient, mounted with Prolong Gold Antifade Mountant (P36930) and DAPI (P36931) if necessary, and sealed with clear nail polish.

### *ACTN4 association* with heart failure with preserved ejection fraction in humans

We assessed for associations between genetic variation in *ACNT4* and human disease phenotypes using BioVU, the Vanderbilt University biobank linking de-identified electronic health records with DNA samples and genotype data^68^. Analyses were conducted in a cohort of 70,854 adults of European ancestry (EA) previously genotyped on the Illumina Multi-Ethnic Genotyping Array (MEGA^EX^). Quality control (QC) analyses used PLINK v 1.90β3 ^95^ and included reconciling strand flips, verifying that allele frequencies were concordant among data sets, and identifying duplicate and related individuals^96^. Data were standardized using the HRC-1000G-check tool v4.2.5 (http://www.well.ox.ac.uk/~wrayner/tools/) and pre-phased using SHAPEIT^97^. Data were imputed using IMPUTE2^98^ in conjunction with the 1000 Genomes cosmopolitan reference haplotypes. In order to evaluate whether *ACTN4* is linked with human clinical phenotypes consonant with pre-clinical data, we tested for association between independent (r^2^<0.05) SNPs in *ACTN4* (within 1 Mb of gene boundary) and the cognate phenotype of heart failure with preserved ejection fraction (HFpEF), defined using a validated computational algorithm. Association testing utilized logistic regression adjusted for age, sex, and the first 10 principal components (PCs) of ancestry. A Benjamini-Hochberg correction was used to account for multiple testing and a false discovery rate (FDR) of <0.1 used for statistical significance.

Independent ACTN4 SNPs used in analyses: rs1985636; rs117885633; rs117784702; rs78871891; rs112979658; rs150193161; rs12981194; rs113293049; rs117124256; rs139207522; rs140381330; rs138414242; rs749700; rs543111965

### Statistical information

**Figure 2: 2A+B**: n=127 si-Scr, 99 si-*ACTN4*, 101 si-*ACTN4* + *ACTN4*-mEGFP cells, Kruskal-Wallis multiple comparisons test; **2C**: n=127 mEGFP, 136 *ACTN4*-mEGFP, 76 mEmerald, 85 *ACTN2*-mEmerald cells, Mann-Whitney test; **2D**: n=105 GFP, 92 *ACTN2*-GFP, 104 *ACTN4*-GFP, 106 *ACTN4*ABD:*ACTN2*-GFP, 92 *ACTN4*K255E-GFP cells, Brown-Forsythe and Welch’s ANOVA multiple comparisons test; **2E+F**: n= 56 si-Scr, 68 si-*ACTN4* cells, unpaired t-test with Welch’s correction; **2F (right):** n=44 *ACTN2-* halo, 36 *ACTN-4*halo cells, unpaired t-test with Welch’s correction; **2G+H:** n= 109 si-Scr, 112 si-*ACTN4* cells, Welch’s t-test.

**Figure 3: 3A+B:** n=98 si-Scr, 83 si-ACTN4 cells (passive), unpaired t-test with Welch’s correction; n=101 si-Scr, 84 si-*ACTN4* cells (active), unpaired t-test with Welch’s correction; **3C+D:** n=9 si-Scr, 13 si-*ACTN*4 cells, unpaired t-test with Welch’s correction; **3E (general hypertrophy):** ACTA1, p=0.06826908 (ns); ACTC1, p=0.01280907 (up); CRYAB, p=0.08078119 (ns); CSRP3, p=0.45501499 (ns); GAPDH, p=0.00210705 (up); MYL3, p=0.02767114 (up); MYL9, p=1.48E-10 (up); SLC25A3, p=0.91314822 (ns); SLC25A4, p=0.00056612 (up); TNNC1, p=0.00016045 (up); TPM1, p=0.00329765 (up); NPPA, p=0.03418045 (up); XIRP2, p=2.57E-40 (up); MYH7, p=0.06316123 (ns); **3E (HCM associated):** CLOCK, p=0.76085429 (ns); CREB1, p=0.01892334 (down); ETV1, p=2.71E-05 (down); FOXN3, p=4.99E-05 (up); MEF2A, p=0.73009667 (ns); MITF, p=0.31718673 (ns); MXI1, p=1.89E-05 (down); NFE2L1, p=0.00012849 (up); REST, p=0.15478464 (ns); SREBF2, p=0.01860801 (up); ZNF91, p=0.95826724 (ns), p-values generated using DESeq2 (25516281); **3F+3G:** n=176 si-Scr, 114 si-*ACTN4*+dmso, 136 si-*ACTN4*+50nM, 142 si-*ACTN4*+100nM, 135 si-*ACTN4*+250nM, 145 si-*ACTN4*+500nM cells, Kruskal-Wallis multiple comparisons test; **3I:** n=19 siScr, 20 si-*ACTN4* cells, Mann-Whitney test; **3J:** n=19 siScr, 20 si-*ACTN4* cells, si-Scr vs. si-*ACTN4* pre-drug vs. pre-drug and post-drug vs. post-drug: Mann-Whitney test; si-Scr pre-vs. post-drug and si-*ACTN4* pre-vs. post-drug: Wilcoxon matched-pairs signed rank test.

**Figure 4: 4B:** n=19 control, 19 *ACTN4-*MO embryos, t-test with Welch’s correction; **4D:** n=53 non-injected control 1, 55 *ACTN4-MO1*-injected, 25 non-injected control 2, 23 *ACTN4-*MO-injected embryos, unpaired t-test with Welch’s correction; **4E:** 57 *ACTN4-MO1*-injected, 23 non-injected control 2, 20 *ACTN4-*MO-injected embryos, unpaired t-test with Welch’s correction.

**Figure 5: 5E-H:** n=10 NIC, 14 *ACTN4*-MO, 13 h*ACTN4*resc, 23 *ACTN4*-MO+mava embryos, Brown-Forsyth and Welch ANOVA; **5I:** n=37 embryos, Pearson’s r; **5J-M:** n=33 NIC, 31 *ACTN4*-MO, 24 h*ACTN4*Resc embryos, Brown-Forsyth and Welch ANOVA.

**Figure 6: 6C-F:** n=15 DMSO, 17 OM embryos, Welch’s t-test.

**Supplemental Figure 2: S2D-E:** n=132 si-Scr, 139 si-*ACTN4* cells, Mann-Whitney test; **S2F:** n=127 si-Scr, 99 si-*ACTN4*, 101 si-*ACTN4* + *ACTN4*-mEGFP cells, Kruskal-Wallis multiple comparisons test; **S2G:** n=55 si-Scr, n=43 si-*ACTN4* cells, Welch’s t-test.

## Acknowledgements

**Acknowledgments** We would like to thank Kari Seedle at the Vanderbilt Nikon Center for Excellence and the Vanderbilt Center for Imaging Shared Resources (CISR) for experimental and imaging assistance. We also thank the Vanderbilt Cardiology Core for graciously providing samples of normal human ventricular myocardium tissue, and the VANTAGE: Vanderbilt Technologies for Advanced Genomics for assistance with RNAseq data processing and analysis.

## Sources of Funding

This work was supported by a grant from the National Institute of General Medical Sciences (R35 GM125028) and an Expansion Award from Additional Ventures (1162039) to DTB and the following graduate student fellowships from the American Heart Association: (836090) to JBH, (1070985) to ZCS, and (18PRE33960551) to NT.

## SUPPLEMENTAL FIGURES

**Figure S1.**
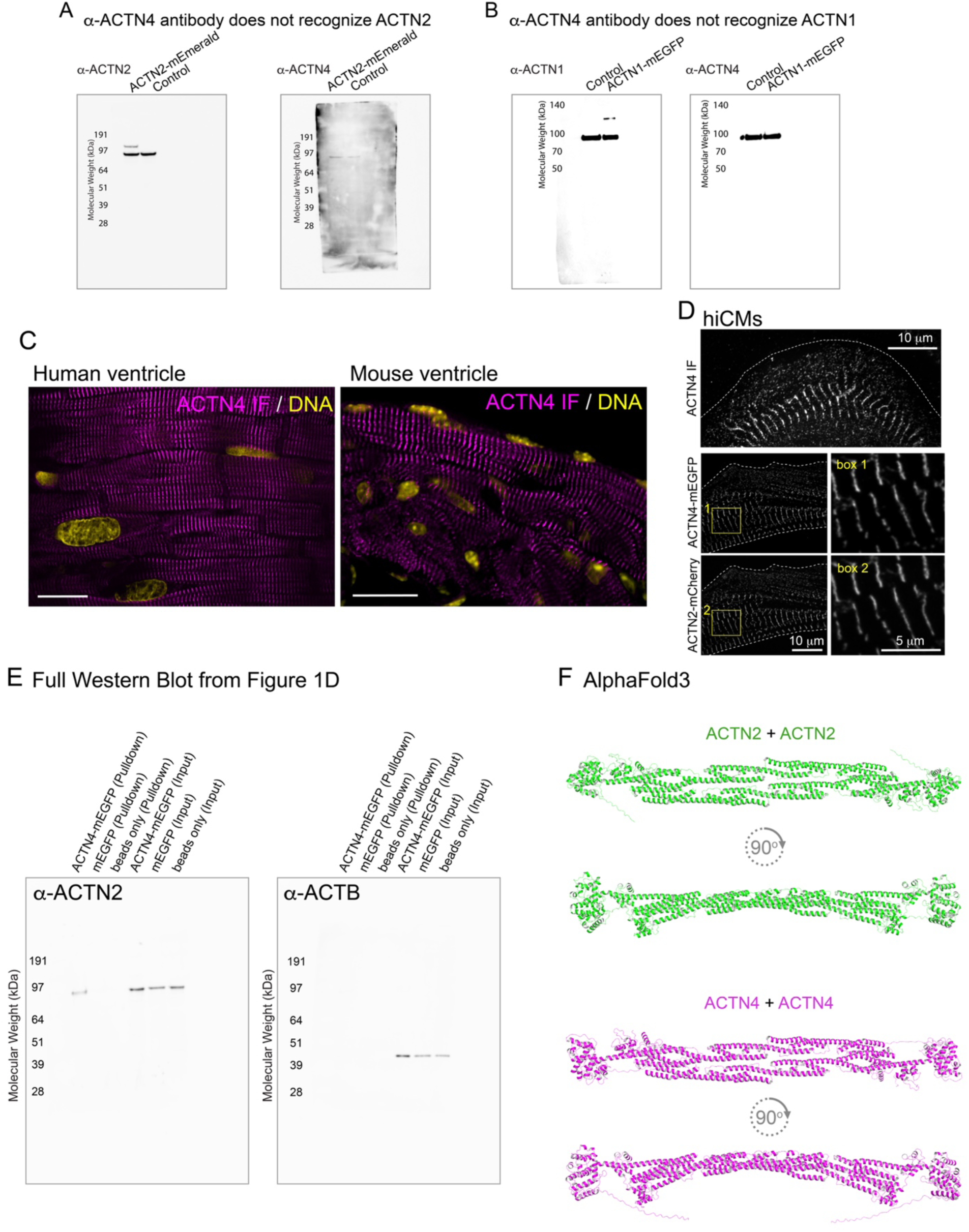
**S1A:** Left panel: western blot of whole cell lysate from hiCMs expressing either ACTN2-mEmerald (left lane) or vehicle (right lane), blotted for ACTN2. Right panel: Blot from left panel stripped and re-blotted using antibodies raised against ACTN4. ACTN4 antibodies do not recognize larger band (representing ACTN2-mEmerald). **S1B:** Left panel: western blot of whole cell lysate from U2OS cells expressing either vehicle (left lane) or ACTN1-mEGFP (right lane). Right panel: blot from left panel stripped and re-blotted using antibodies raised against ACTN4. ACTN4 antibodies do not recognize larger band (representing ACTN1-mEGFP). **S1C:** Left panel: ACTN4 (magenta) and DNA (yellow) stained in a formalin-fixed paraffin-embedded (FFPE) tissue section of the adult human heart ventricle. Right panel: ACTN4 (magenta) and DNA (yellow) stained in a formalin-fixed paraffin-embedded (FFPE) tissue section of the adult mouse heart ventricle. **S1D:** Top: ACTN4 stained in a single hiCM. Bottom: ACTN4-mEGFP and ACTN2-mEGFP co-expressed within a hiCM. Boxes show higher mag inlays. **S1E:** Left: blotted with antibodies against ACTN2. Right: blotted with antibodies against actin (ACTB). **S1F:** Top: AlphaFold3-predicted structure of an ACTN2 homodimer. Bottom: AlphaFold3-predicted structure of an ACTN4 homodimer.

**Figure S2.**
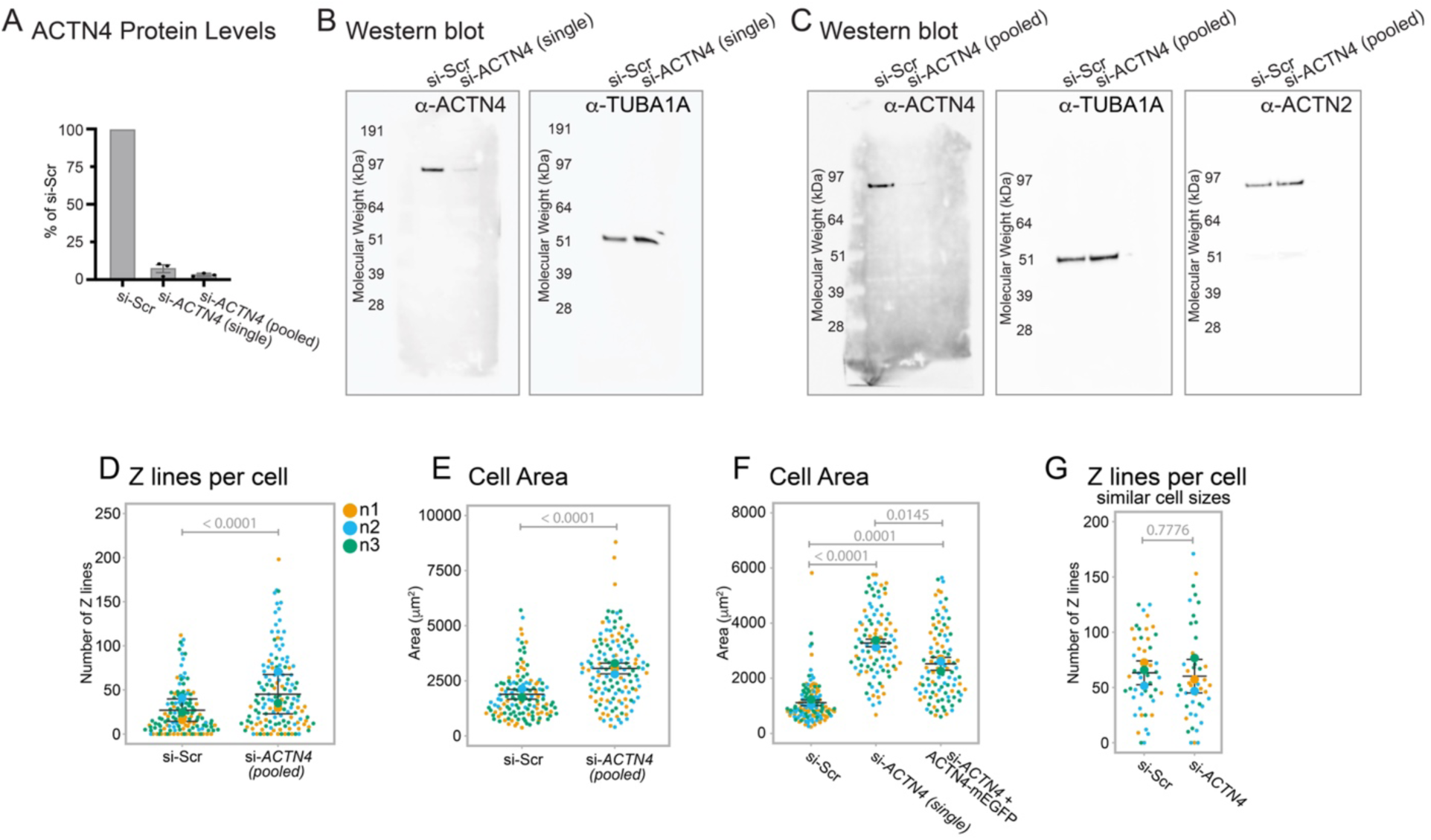
**S2A:** ACTN4 protein levels (as a % of control) following depletion of ACTN4 using two different siRNAs. **S2B:** Left: representative western blot of ACTN4 in hiCMs following treatment with either non-targeting (si-Scr) or *ACTN4-*targeting (si-*ACTN4*) single siRNA. Middle: Same blot as left stripped and blotted for tubulin (TUB1A, used as loading control). **S2C:** Left: representative western blot of ACTN4 in hiCMs following treatment with either non-targeting (si-Scr) or pooled *ACTN4-*targeting (si-*ACTN4*) siRNA. Middle: Same blot as left stripped and blotted for tubulin (TUB1A, used as loading control). Right: Same blot as left two blots stripped and blotted for ACTN2. **S2D:** Z-lines per cell quantified in hiCMs treated with either non-targeting (si-Scr) or pooled *ACTN4-*targeting (si-*ACTN4*) siRNA. **S2E:** Cell area of hiCMs treated with either non-targeting (si-Scr) or pooled *ACTN4-*targeting (si-*ACTN4*) siRNA. **S2F:** Cell area of hiCMs treated with either non-targeting (si-Scr), single *ACTN4-*targeting (si-*ACTN4*), or si-*ACTN4* + expressing ACTN4-mEGFP. **S2G:** Z-lines per cell in hiCMs following treatment with either non-targeting (si-Scr) or *ACTN4-*targeting (si-*ACTN4*) siRNA. Only hiCMs with area between 1500-3000um^2 included in analysis.

**Figure S3.**
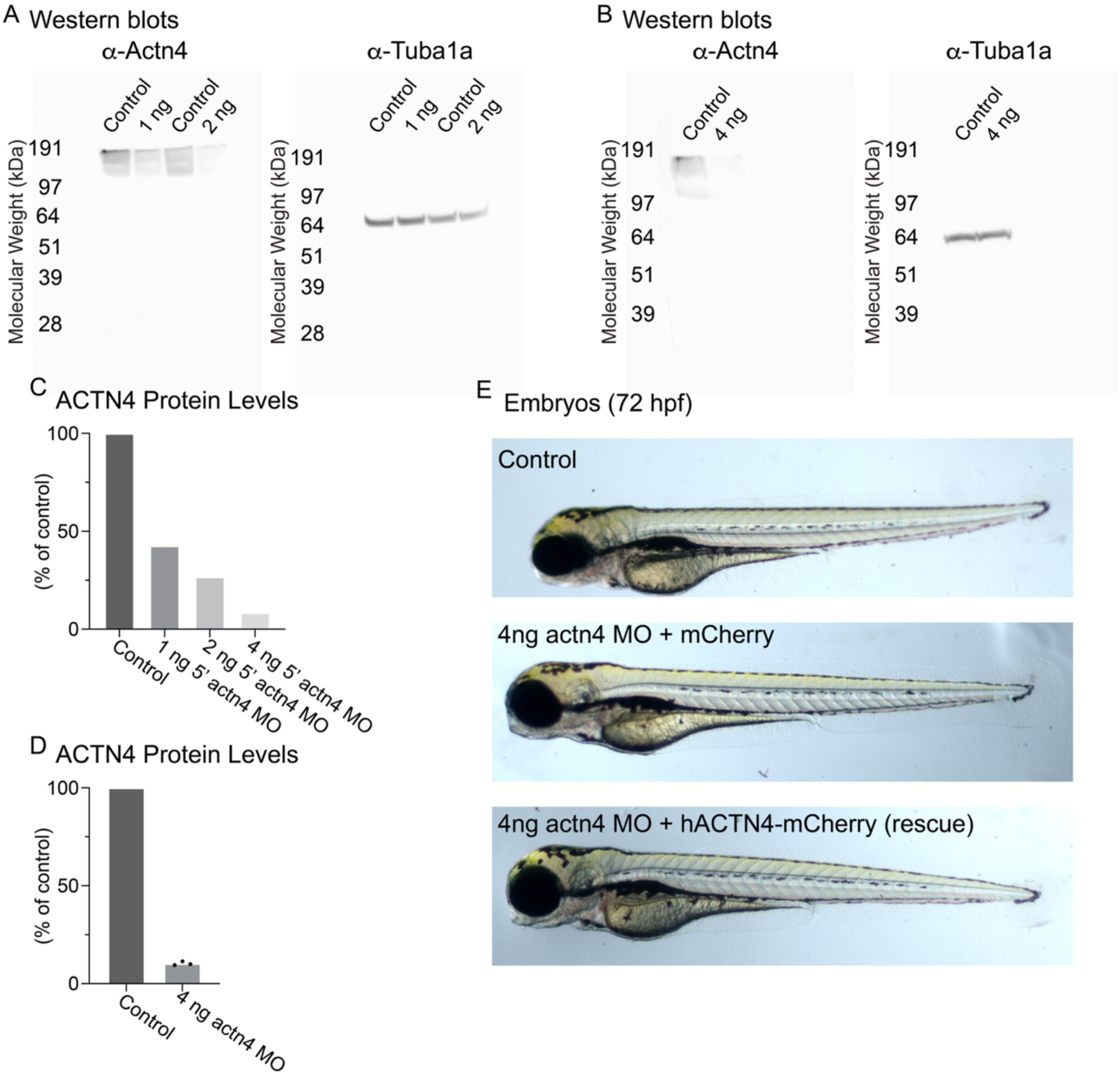
**S3A-B:** western blots showing actn4 levels in whole zebrafish lysate following either no injection (control) or injection with 1ng, 2ng, or 4ng 5’-UTR-targeting actn4 morpholinos. Anti-actn4 shows actn4 blots and anti-tub1a shows tubulin blots used as loading control. **S3C-D:** quantification of western blots showing actn4 protein after morpholino injections compared to control (non-injected) embryos at 72hpf. **S3E:** representative images of 72hpf zebrafish embryos that were either non-injected (control) or injected at one-cell stage with 4ng *actn4-*targeting morpholinos (*actn4* MO) or 4ng *actn4-*targeting morpholinos and mRNA encoding human ACTN4-mCherry (*actn4* MO + hACTN4-mCherry).

## Notes

### Competing Interest Statement

The authors have declared no competing interest.

